# Gene co-expression network analysis reveal core responsive genes in *Parascaris univalens* tissues following ivermectin exposure

**DOI:** 10.1101/2023.12.07.570202

**Authors:** Faruk Dube, Nicolas Delhomme, Frida Martin, Andrea Hinas, Magnus Åbrink, Staffan Svärd, Eva Tydén

## Abstract

Anthelmintic resistance in equine parasite *Parascaris univalens*, compromises ivermectin (IVM) effectiveness and necessitates an in-depth understanding of its resistance mechanisms. Most research, primarily focused on holistic gene expression analyses, may overlook vital tissue-specific responses and often limit the scope of novel genes. This study leveraged gene co-expression network analysis to elucidate tissue-specific transcriptional responses and to identify core genes implicated in the IVM response in *P. univalens*. Adult worms (n=28) were exposed to 10^-11^ M and 10^-9^ M IVM *in vitro* for 24 hours. RNA-sequencing examined transcriptional changes in the anterior end and intestine. Differential expression analysis revealed pronounced tissue differences, with the intestine exhibiting substantially more IVM-induced transcriptional activity. Gene co-expression network analysis identified seven modules significantly associated with the response to IVM. Within these, 219 core genes were detected, largely expressed in the intestinal tissue and spanning diverse biological processes with unspecific patterns. After 10^-11^ M IVM, intestinal tissue core genes showed transcriptional suppression, cell cycle inhibition, and ribosomal alterations. Interestingly, genes *PgR028_g047* (*sorb-1*), *PgB01_g200* (*gmap-1*) and *PgR046_g017* (*col-37* & *col-102*) switched from downregulation at 10^-11^ M to upregulation at 10^-9^ M IVM. The 10^-9^ M concentration induced expression of cuticle and membrane integrity core genes in the intestinal tissue. No clear core gene patterns were visible in the anterior end after 10^-11^ M IVM. However, after 10^-9^ M IVM, the anterior end mostly displayed downregulation, indicating disrupted transcriptional regulation. One interesting finding was the non-modular calcium-signaling gene, *PgR047_g066 (gegf-1)*, which uniquely connected 71 genes across four modules. These genes were enriched for transmembrane signaling activity, suggesting that *PgR047_g066 (gegf-1)* could have a key signaling role. By unveiling tissue-specific expression patterns and highlighting biological processes through unbiased core gene detection, this study reveals intricate IVM responses in *P. univalens*. These findings suggest alternative drug uptake of IVM and can guide functional validations to further IVM resistance mechanism understanding.

**Author summary:** In our study, we tackled the challenge of understanding how the equine roundworm *Parascaris univalens* has become resistant to ivermectin (IVM). We exposed adult worms in laboratory conditions to IVM and thereafter dissected two tissues, the frontal part and the intestine of the worm. We used gene networks and focused on how these two tissues respond at the genetic level to exposure of IVM. We discovered that the response to IVM is highly tissue-specific. The intestinal tissue, in particular, showed a much stronger reaction to the drug compared to the frontal part of the worm. We identified 219 key genes, mainly in the intestinal tissue, involved in various biological functions that play a crucial role in how the parasite deals with IVM. Interestingly, we found a decrease in gene activity leading to cellular disruptions at lower drug concentration, whereas genes responsible for maintaining the worm’s structural integrity were triggered at high concentration. One of our significant finding was the identification of, *PgR047_g066 (gegf-1)*, which seems to act as a master regulator, coordinating the response of numerous other genes. This finding opens new avenues for understanding the complex ways in which *P. univalens* respond to drug treatment. Our research not only sheds light on the specific ways *P. univalens* responds to IVM, but it also demonstrates the power of looking at gene networks to uncover new and important genes. These insights can be crucial for developing new strategies to combat drug resistance in parasites, a matter of great importance in both veterinary and human medicine.

## Introduction

Parasitic nematodes represent a significant health challenge, causing persistent and debilitating diseases in humans and animals [1, 2]. The primary strategy to control parasitic nematodes relies on treatment with anthelmintic drugs such as ivermectin (IVM), which belongs to the Macrocyclic lactones (MLs) drug class. This drug class acts by binding to parasite specific glutamate-gated ion channels (GluCls) and γ-aminobutyric acid (GABA) receptors, resulting in pharyngeal hyperpolarization and flaccid muscle paralysis [3–5]. Unregulated preventive treatment and overuse of anthelmintic drugs has contributed to the development of resistance in several parasites of veterinary importance [6]. One such is *Parascaris* spp. an equine parasitic nematode prevalent in foals and yearlings. Infection manifestations of this gastrointestinal nematode mainly includes nasal discharge, coughing and impaired growth, while large burdens can be lethal due to obstruction and perforation of the small intestine [7, 8]. The first reported case of anthelmintic resistance in *Parascaris* spp. was to IVM in 2002 [9] and since then resistance has been reported around the world and is now considered wide spread [10]. The problem with drug resistance, together with the risk of lethal complications in infected foals makes *Parascaris* spp. a major threat to the equine industry and equine health.

While the specific mechanisms underlying IVM resistance in *Parascaris* spp. remain elusive, our limited understanding is largely derived from studies in the free-living nematode, *Caenorhabditis elegans*, and a few other parasitic nematodes [11]. Generally, resistance mechanisms are proposed to be due to mutations or differential expression in genes encoding drug targets, metabolic enzymes, transporters and key transcription factors [11–14]. For example, mutations in the GluCl gene, *avr-14* has been reported to confer ML resistance in *C. elegans* [15], *Cooperia oncophora* [16] and *Haemonchus contortus* [17]. In addition, decreased expression of the GluCl genes, *glc-3* and *glc-5* has been implicated in ML resistance in *H. contortus* [18]. Furthermore, increase dexpression of transporter genes of the ATP-binding cassette family, particularly P-glycoproteins (Pgps), also known as efflux pumps have been observed in nematodes with an ML-resistant phenotype [19, 20]. These efflux proteins function to expel drugs from the cell, thereby preventing them from reaching their target sites. However, increased expression of Pgps was not been observed in *Parascaris* spp. following IVM exposure [21, 22]. Instead, other transport families such as the major facilitator superfamily and the solute carrier superfamily were found to be differentially expressed in IVM exposed *Parascaris* spp. worms [22]. Collectively, this highlights the likely multigenic basis of anthelmintic resistance in Ascarids/*Parascaris* spp.

Upon oral administration in horses, IVM reaches high concentrations in the small intestine, where *Parascaris* spp. adults reside within the lumen [23]. Given the phylogenetic proximity of *Parascaris* spp. to *Ascaris suum* within the Ascarididae order [24], insights from *A. suum* suggest that IVM is mainly absorbed through transcuticular diffusion [25]. Consequently, it is likely that IVM also penetrates the *Parascaris* spp. cuticle by passive diffusion, leading to its accumulation within the parasite but ingestion of semi-digested intestinal contents can potentially serve as a supplementary route of exposure. The nematode intestine is a key organ that interacts with the environment, plays an active role in nutrient digestion, cellular trafficking, metabolic pathways, and defense against xenobiotics, including drugs [26, 27]. Thus, the intestinal tissue is a pivotal area of study in understanding responses to drug exposure, potentially providing insights into resistance mechanisms. Despite this, few studies have explored differential gene expression in the intestinal tissue of parasitic nematodes post-drug exposure. The large size of adult *Parascaris* spp., however, makes such investigations feasible.

Another important region of nematodes is the anterior that houses essential structures, such as the pharynx, ganglia, and nerve ring, which play significant roles in the nematode’s biology, including food intake, neural function, and sensory processing [28]. Ivermectin is known to interact with the nervous system [3–5], and changes in gene expression in these areas could provide insights into how the drug affects these functions. Therefore, analyzing gene expression responses in the anterior region and intestine after IVM exposure may reveal tissue-specific differences in drug response and resistance mechanisms, providing insights into the overall parasite’s response and resistance.

The advent of omics technologies and enhanced computational capabilities, underscored by the recent publication of a draft genome for *Parascaris univalens* [29], have led to new avenues for exploring potential drug targets and resistance mechanisms. Conventional candidate gene approaches [13, 22, 30–33], often biased by selectively choosing candidates from existing literature or based solely on the expression levels, are constrained by their reliance on pre-existing knowledge, limiting the discovery of novel candidates. In contrast, the rise of data-driven analytical methods, including both supervised and unsupervised techniques [34–36], such as those used in gene network building and analysis [37, 38], facilitate a more objective exploration of gene functions and interactions. These methods provide insights into the multifaceted gene relationships that govern drug response and/or resistance, without the predispositions inherent in conventional approaches. For instance, the Seidr toolkit [37], a paradigm of such unbiased methodologies, was recently utilized to construct a gene network for Norway spruce (*Picea abies*) under drought conditions, successfully pinpointing key genes and inferring functions for uncharacterized ones [37]. This example illustrates Seidr’s capacity for conducting comprehensive gene network analyses, which can be instrumental in deciphering complex biological phenomena, including drug responses and resistance mechanisms in parasitic nematodes.

Considering the risk of lethal complications in foals infected with *Parascaris* spp. and the lack of new anthelmintic drugs for the equine market, it is crucial to gain understanding of the underlying causes of resistance in this parasite species. In this study, we undertook a comprehensive examination of gene expression within two vital anatomical regions of *P. univalens* adult worms exposed *in vitro* to IVM at concentrations of 10^-9^ M and 10^-11^ M for 24 hours. Leveraging the Seidr toolkit [37], we adopted a less biased, data-driven gene network approach to systematically identify and evaluate the core genes underlying IVM response and potential resistance mechanisms in the nerve-dominated anterior region and the metabolically active intestinal tissue.

## Results

### Sequencing depth and quality control

The sequencing process yielded 23-35 million reads per sample from batch A (S1 flow cell) and 66-135 million reads per sample from batch B (S4 flow cell). After quality control with SortMeRNA and Trimmomatic, we obtained 20-33 and 58-124 million reads per sample for sequencing batch A [22] and B, respectively. For both batches, about 90% of all the quality controlled reads mapped to the reference transcriptome (see Materials and methods).

### Tissue-specific variability in differentially expressed genes after ivermectin exposure

After exposure to IVM, a notable disparity in gene expression changes was observed between the intestinal tissue and anterior end. The intestinal tissue displayed a substantial response, with 2465 DEGs after IVM exposure. In contrast, the anterior end exhibited 11-fold fewer DEGs, revealing only 227 genes (**S1 Table**). A set of 38 DEGs were shared between tissue types and IVM concentrations (**Table 1**).

**Table 1.**
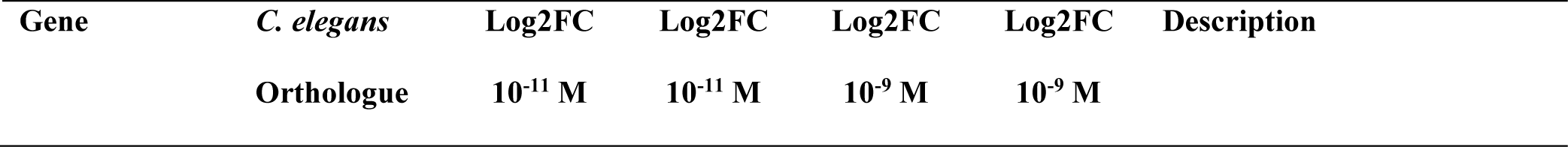
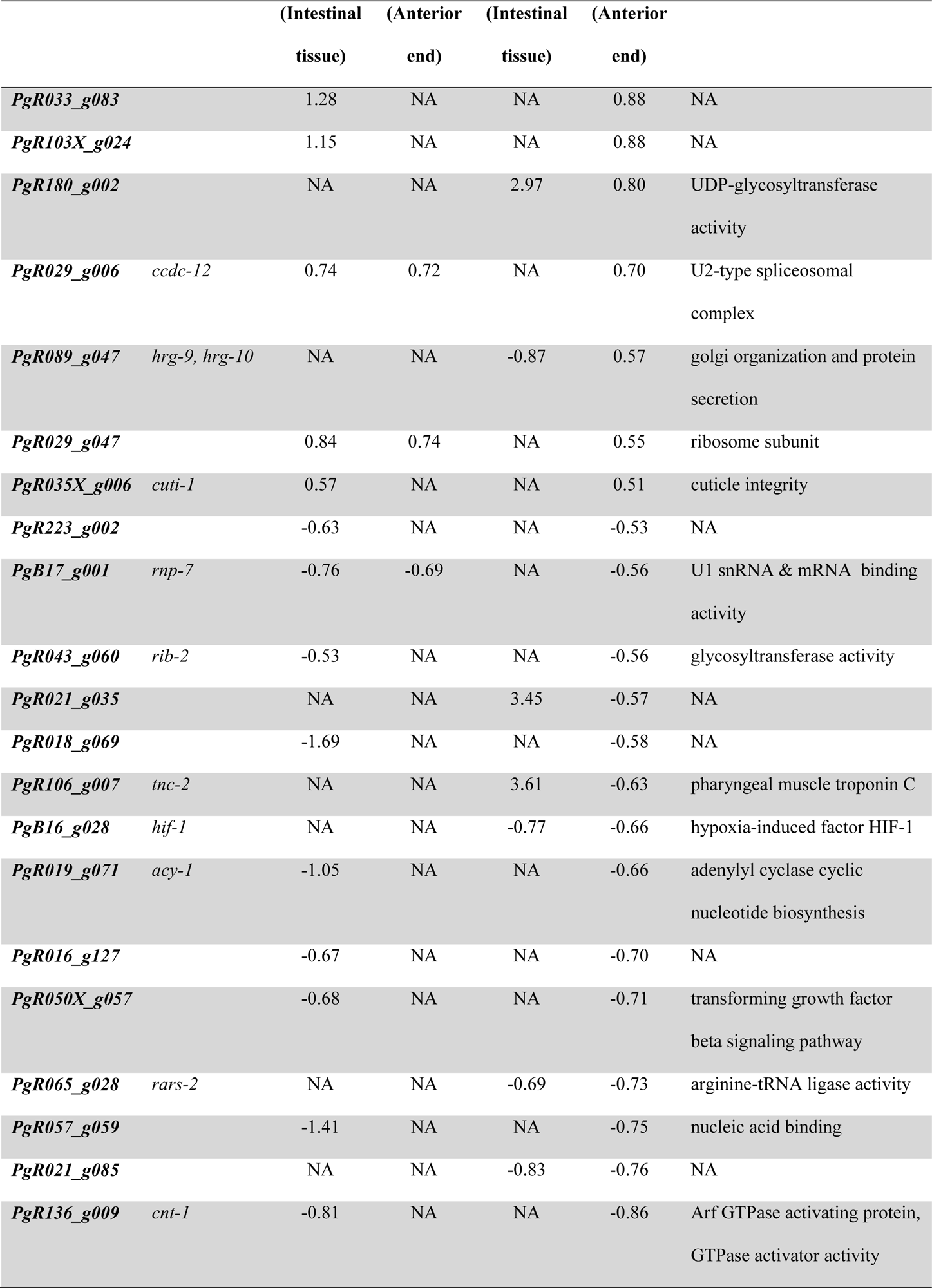
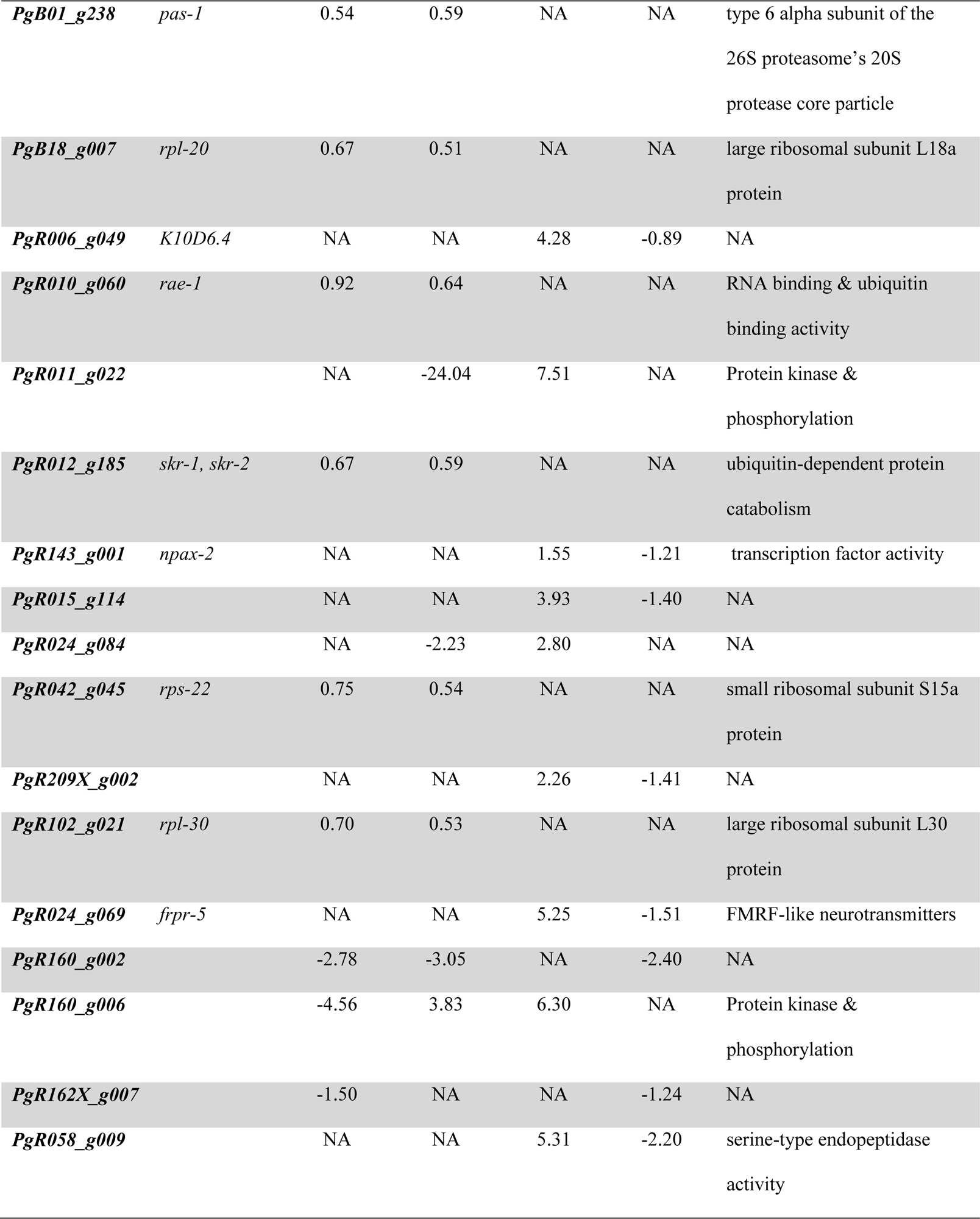
Differentially expressed genes common between intestinal tissue and anterior end for worms exposed to 10^-11^ M and 10^-9^ M ivermectin.

| Gene | *C. elegans* | Log2FC | Log2FC | Log2FC | Log2FC | Description |
| --- | --- | --- | --- | --- | --- | --- |
| | Orthologue | $10^{-11}$ M | $10^{-11}$ M | $10^{-9}$ M | $10^{-9}$ M | |

|  |  | (Intestinal<br>tissue) | (Anterior<br>end) | (Intestinal<br>tissue) | (Anterior<br>end) |  |
| --- | --- | --- | --- | --- | --- | --- |
| **PgR033_g083** |  | 1.28 | NA | NA | 0.88 | NA |
| **PgR103X_g024** |  | 1.15 | NA | NA | 0.88 | NA |
| **PgR180_g002** |  | NA | NA | 2.97 | 0.80 | UDP-glycosyltransferase<br>activity |
| **PgR029_g006** | *ccdc-12* | 0.74 | 0.72 | NA | 0.70 | U2-type spliceosomal<br>complex |
| **PgR089_g047** | *hrg-9, hrg-10* | NA | NA | -0.87 | 0.57 | golgi organization and protein<br>secretion |
| **PgR029_g047** |  | 0.84 | 0.74 | NA | 0.55 | ribosome subunit |
| **PgR035X_g006** | *cuti-1* | 0.57 | NA | NA | 0.51 | cuticle integrity |
| **PgR223_g002** |  | -0.63 | NA | NA | -0.53 | NA |
| **PgB17_g001** | *rnp-7* | -0.76 | -0.69 | NA | -0.56 | U1 snRNA & mRNA binding<br>activity |
| **PgR043_g060** | *rib-2* | -0.53 | NA | NA | -0.56 | glycosyltransferase activity |
| **PgR021_g035** |  | NA | NA | 3.45 | -0.57 | NA |
| **PgR018_g069** |  | -1.69 | NA | NA | -0.58 | NA |
| **PgR106_g007** | *tnc-2* | NA | NA | 3.61 | -0.63 | pharyngeal muscle troponin C |
| **PgB16_g028** | *hif-1* | NA | NA | -0.77 | -0.66 | hypoxia-induced factor HIF-1 |
| **PgR019_g071** | *acy-1* | -1.05 | NA | NA | -0.66 | adenylyl cyclase cyclic<br>nucleotide biosynthesis |
| **PgR016_g127** |  | -0.67 | NA | NA | -0.70 | NA |
| **PgR050X_g057** |  | -0.68 | NA | NA | -0.71 | transforming growth factor<br>beta signaling pathway |
| **PgR065_g028** | *rars-2* | NA | NA | -0.69 | -0.73 | arginine-tRNA ligase activity |
| **PgR057_g059** |  | -1.41 | NA | NA | -0.75 | nucleic acid binding |
| **PgR021_g085** |  | NA | NA | -0.83 | -0.76 | NA |
| **PgR136_g009** | *cnt-1* | -0.81 | NA | NA | -0.86 | Arf GTPase activating protein,<br>GTPase activator activity |
| ***PgB01_g238*** | *pas-1* | 0.54 | 0.59 | NA | NA | type 6 alpha subunit of the<br>26S proteasome's 20S<br>protease core particle |
| ***PgB18_g007*** | *rpl-20* | 0.67 | 0.51 | NA | NA | large ribosomal subunit L18a<br>protein |
| ***PgR006_g049*** | *K10D6.4* | NA | NA | 4.28 | -0.89 | NA |
| ***PgR010_g060*** | *rae-1* | 0.92 | 0.64 | NA | NA | RNA binding & ubiquitin<br>binding activity |
| ***PgR011_g022*** |  | NA | -24.04 | 7.51 | NA | Protein kinase &<br>phosphorylation |
| ***PgR012_g185*** | *skr-1, skr-2* | 0.67 | 0.59 | NA | NA | ubiquitin-dependent protein<br>catabolism |
| ***PgR143_g001*** | *npax-2* | NA | NA | 1.55 | -1.21 | transcription factor activity |
| ***PgR015_g114*** |  | NA | NA | 3.93 | -1.40 | NA |
| ***PgR024_g084*** |  | NA | -2.23 | 2.80 | NA | NA |
| ***PgR042_g045*** | *rps-22* | 0.75 | 0.54 | NA | NA | small ribosomal subunit S15a<br>protein |
| ***PgR209X_g002*** |  | NA | NA | 2.26 | -1.41 | NA |
| ***PgR102_g021*** | *rpl-30* | 0.70 | 0.53 | NA | NA | large ribosomal subunit L30<br>protein |
| ***PgR024_g069*** | *frpr-5* | NA | NA | 5.25 | -1.51 | FMRF-like neurotransmitters |
| ***PgR160_g002*** |  | -2.78 | -3.05 | NA | -2.40 | NA |
| ***PgR160_g006*** |  | -4.56 | 3.83 | 6.30 | NA | Protein kinase &<br>phosphorylation |
| ***PgR162X_g007*** |  | -1.50 | NA | NA | -1.24 | NA |
| ***PgR058_g009*** |  | NA | NA | 5.31 | -2.20 | serine-type endopeptidase<br>activity |

After exposure of worms to the 10^-11^ M IVM concentration, the variance in gene expression responses between the two tissue types was even more pronounced. The intestinal tissue revealed 1287 DEGs, whereas the anterior end had only 57, resulting in a 23-fold difference (**S1 Table**). Among these, 11 DEGs were common to both tissues. Notably, all shared DEGs followed a similar expression pattern in both tissue types except for *PgR160_g006*, which displayed an opposite trend. These shared DEGs are generally implicated in mRNA processing, translation, and protein catabolism (**Table 2**). Overall, a downregulated profile was observed for most DEGs in the intestinal tissue. As many as 58% (758 genes) of DEGs were downregulated, with 38% (491 genes) exceeding lowered expression. Only 14% (179 genes) of the intestinal tissue DEGs exhibited an upregulation above 2-fold. In contrast, the anterior end showed an upregulated profile of DEGs with 66% (45 genes) being upregulated of which 35% (24 genes) surpassed a 2-fold change, and 24% (16 genes) displayed a downregulation greater than 2-fold (**S1 Table**).

**Table 2.** Differentially expressed genes common between intestinal tissue and anterior end for worms exposed to 10^-11^ M ivermectin.

| Gene | *C. elegans* | Log2FC | Log2FC | Description |
| --- | --- | --- | --- | --- |
| | Orthologue | $10^{-11}$ M | $10^{-11}$ M | |
|  |  | (Intestinal | (Anterior |  |
|  |  | tissue) | end) |  |
| *PgR029_g006* | *ccdc-12* | 0.74 | 0.72 | U2-type spliceosomal complex |
| *PgR029_g047* |  | 0.84 | 0.74 | ribosome subunit |
| *PgB17_g001* | *rnp-7* | -0.76 | -0.69 | U1 snRNA & mRNA binding activity |
| *PgB01_g238* | *pas-1* | 0.54 | 0.59 | type 6 alpha subunit of the 26S proteasome's 20S protease core particle |
| *PgB18_g007* | *rpl-20* | 0.67 | 0.51 | large ribosomal subunit L18a protein |
| *PgR010_g060* | *rae-1* | 0.92 | 0.64 | RNA binding & ubiquitin<br>binding activity |
| *PgR012_g185* | *skr-1, skr-2* | 0.67 | 0.59 | ubiquitin-dependent protein<br>catabolism |
| *PgR042_g045* | *rps-22* | 0.75 | 0.54 | small ribosomal subunit<br>S15a protein |
| *PgR102_g021* | *rpl-30* | 0.70 | 0.53 | large ribosomal subunit L30<br>protein |
| *PgR160_g002* |  | -2.78 | -3.05 | NA |
| *PgR160_g006* |  | -4.56 | 3.83 | protein kinase &<br>phosphorylation |

From worms exposed to the 10^-9^ M IVM concentration, the intestinal tissue maintained its pronounced response, exhibiting 1402 DEGs, in comparison to the more subdued response from the anterior end, with 191 DEGs, an approximate 7-fold difference (**S1 Table**). Notably, there were 13 DEGs common to both tissues (**Table 3**), with the majority showing opposite expression trends, except for *PgR180_g002*, *PgB16_g028 (hif-1)*, and *PgR065_g028* (*rars-2*). These shared DEGs are associated with diverse biological processes. In this concentration, a contrasting pattern was observed with an overall upregulated response compared to IVM 10^-11^ M in the intestinal tissue. A considerable proportion, 84% (1189 genes) of intestinal DEGs were upregulated, with 79% (1120 genes) above a 2-fold change, and only 7% (98 genes) showed downregulation above 2-fold. By contrast, the anterior end showed 59% (121 genes) downregulated profile of DEGs, 11% (23 genes) of which exceeded a 2-fold change, and 7% (15 genes) displayed upregulation above 2-fold (**S1 Table**).

**Table 3.** Differentially expressed genes common between the intestinal tissue and anterior end for worms exposed to 10^-9^ M ivermectin.

| Gene | *C. elegans* | Log2FC | Log2FC | Description |
| --- | --- | --- | --- | --- |
|  | Orthologue | 10<sup>-9</sup> M | 10<sup>-9</sup> M |  |
|  |  | (Intestinal | (Anterior |  |
|  |  | tissue) | end) |  |
| **PgR180_g002** |  | 2.97 | 0.80 | UDP-glycosyltransferase activity |
| **PgR089_g047** | *hrg-9, hrg-10* | -0.87 | 0.57 | Golgi organization and protein secretion |
| **PgR021_g035** |  | 3.45 | -0.57 | NA |
| **PgR106_g007** | *tnc-2* | 3.61 | -0.63 | pharyngeal muscle troponin C |
| **PgB16_g028** | *hif-1* | -0.77 | -0.66 | hypoxia-induced factor HIF-1 |
| **PgR065_g028** | *rars-2* | -0.69 | -0.73 | arginine-tRNA ligase activity |
| **PgR021_g085** |  | -0.83 | -0.76 | NA |
| **PgR006_g049** | *K10D6.4* | 4.28 | -0.89 | NA |
| **PgR143_g001** | *npax-2* | 1.55 | -1.21 | transcription factor activity |
| **PgR015_g114** |  | 3.93 | -1.40 | NA |
| **PgR209X_g002** |  | 2.26 | -1.41 | NA |
| **PgR024_g069** | *frpr-5* | 5.25 | -1.51 | FMRF-like neurotransmitter |
| **PgR058_g009** |  | 5.31 | -2.20 | serine-type endopeptidase activity |

### Enriched biological processes of differentially expressed genes in the intestinal tissue after ivermectin exposure

Over-representation analysis of the DEGs revealed varying responses in the intestinal tissue depending on the IVM concentration. In 10^-11^ M concentration, downregulated genes were associated with the regulation of cellular processes and protein binding (**Fig 1A & S2A Table**). In contrast, upregulated genes were significantly involved in peptide metabolic processes, and activities related to mitochondrial and ribosomal functions (**Fig 1B & S2B Table**). However, after 10^-9^ M IVM exposure, the ORA of upregulated genes highlighted a broad spectrum of enriched biological processes (**S3 Table**). The predominant processes were associated with monoatomic ion transport, maintenance of cuticle integrity, regulation of transmembrane signaling pathways, and activities concerning the cell periphery **(Fig 2**). No significant enriched biological processes were detected for the downregulated genes.

**Fig 1.**
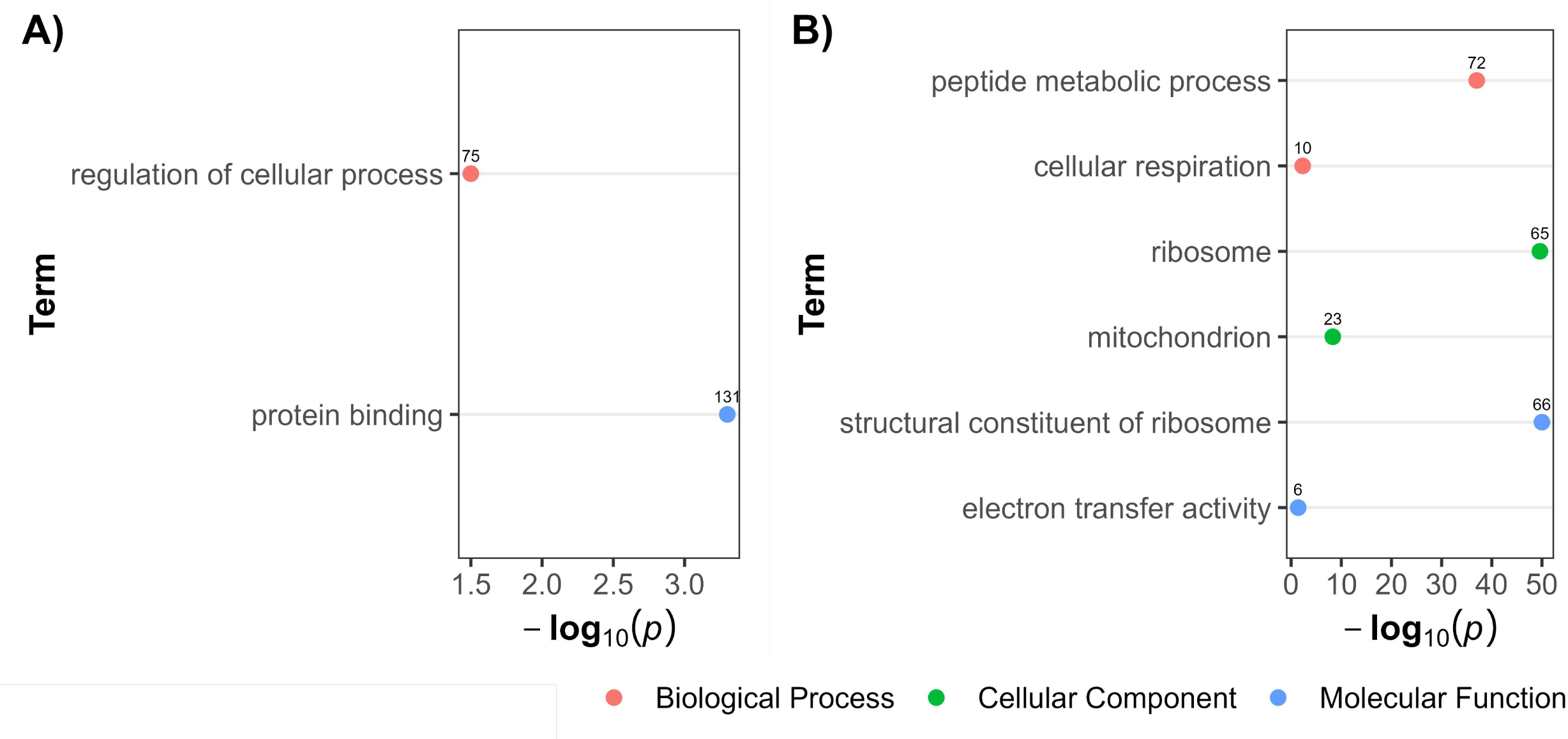
Over-representation analysis showing gene ontology terms in differentially expressed genes in intestinal tissue after worm exposure to 10^-11^ M ivermectin. Panel A presents terms linked to downregulated genes, while Panel B highlights those associated with upregulated genes. The data points, color-coded by ontology classification-Biological Process (red), Cellular Component (green), and Molecular Function (blue) are plotted against their respective -log10 transformed p-values, emphasizing the significance of each term. The numerical labels adjacent to each data point indicate the count of differentially expressed genes associated with the respective GO term.

**Fig 2.**
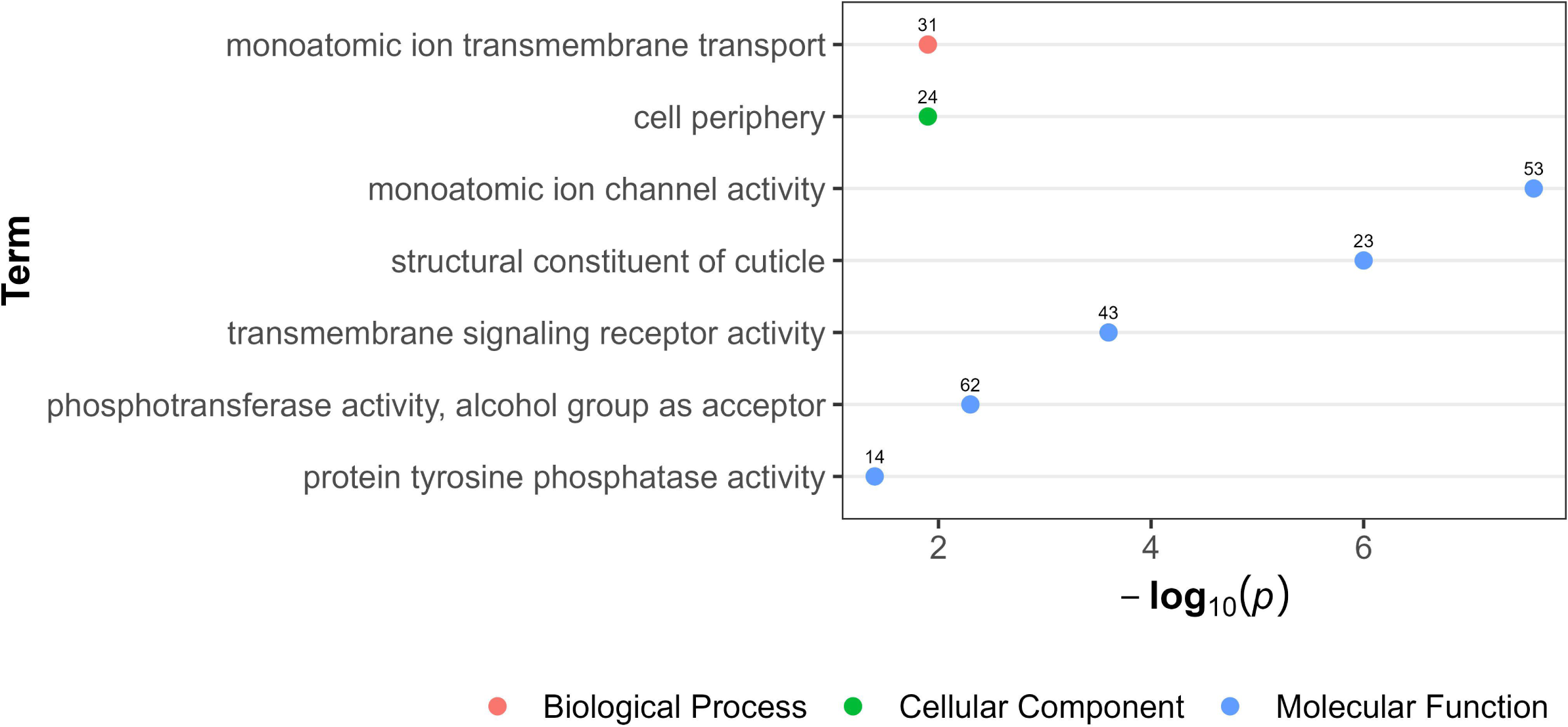
Over-representation analysis showing gene ontology terms in upregulated genes in intestinal tissue after worm exposure to 10^-9^ M ivermectin. The data points, color-coded by ontology classification-Biological Process (red), Cellular Component (green), and Molecular Function (blue) are plotted against their respective -log10 transformed p-values, emphasizing the significance of each term. The numerical labels adjacent to each data point indicate the count of upregulated genes.

### Enriched biological processes of differentially expressed genes in the anterior end after ivermectin exposure

There were no significantly enriched biological processes of the anterior end DEGs in the 10^-11^ M concentration of IVM. The ORA of upregulated genes after 10^-9^ M IVM exposure mainly pointed to involvement in energy-dependent proton transmembrane transportation, which differ compared to the response in the intestinal tissue **(Fig 3 & S4 Table**). No significant enriched biological processes were detected for the downregulated genes.

**Fig 3.**
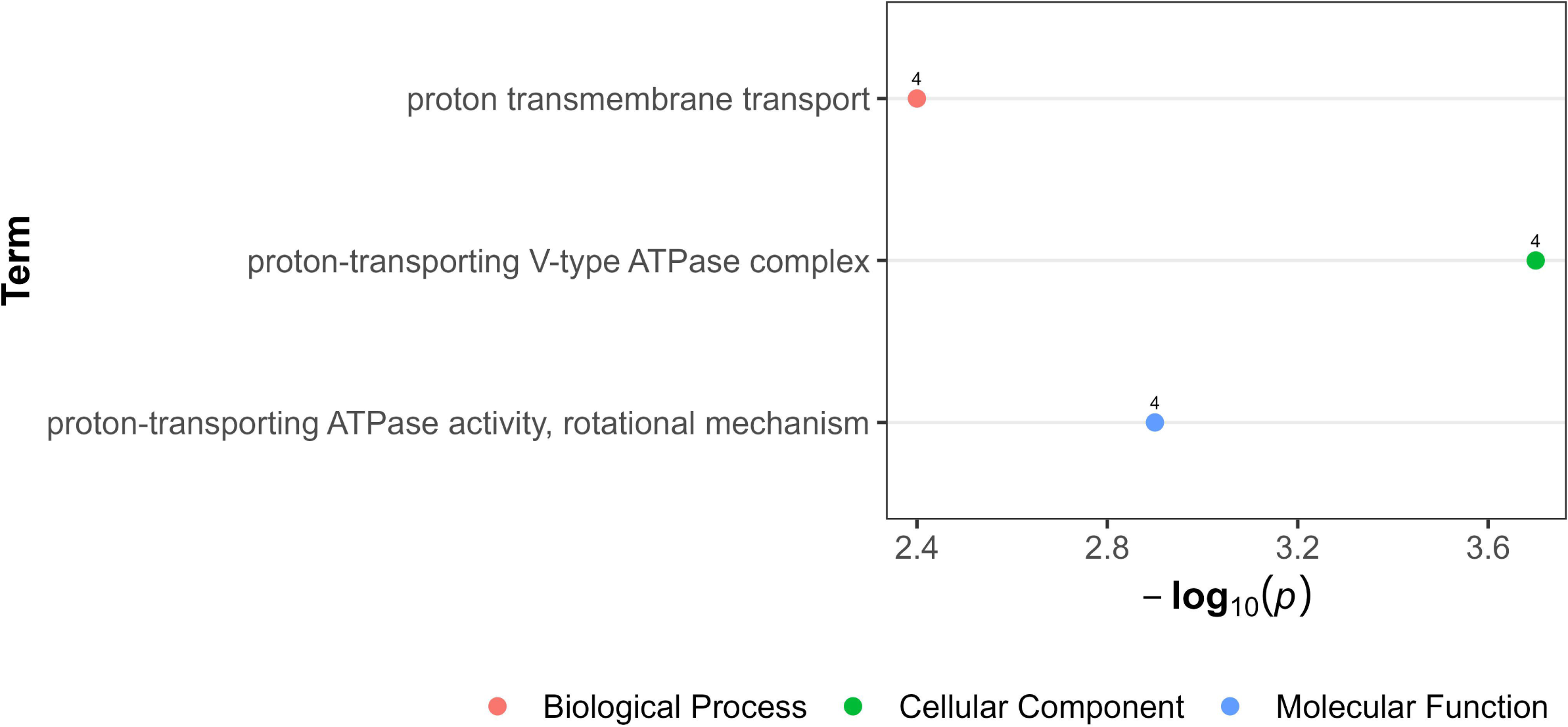
Over-representation analysis showing gene ontology terms in upregulated genes in anterior end after worm exposure to 10^-9^ M ivermectin. The data points, color-coded by ontology classification-Biological Process (red), Cellular Component (green), and Molecular Function (blue) are plotted against their respective -log10 transformed p-values, emphasizing the significance of each term. The numerical labels adjacent to each data point indicate the count of upregulated genes.

### Network analysis reveals gene modules significantly associated with ivermectin response

The application of the Leiden community detection algorithm to partition the consensus network resulted in the identification of 10 discrete gene modules, each characterized by a high correlation in expression profiles. Within this framework, 172 genes were not associated with any module, categorizing them as “non-modular genes”. Notably, two of these genes, *PgR073_g043* and *PgR047_g066 (gegf-1)*, served as connectors between at least two modules, thus being classified as “intermodular genes”. The modules varied in size, containing 788, 811, 1675, 4781, 332, 719, 99, 1147, 104, and 475 genes for Modules 1 through 10, respectively. By conducting overlap statistics of DEGs across tissue types and concentrations with all existing modules, significant associations with the IVM response were identified in seven modules (Modules 1, 2, 4, 5, 7, 8, and 10) (**S1 Fig & S5 Table**).

### Over-representation analysis highlights functional heterogeneity across significant network modules

Subsequent ORA performed on all genes within the significant modules revealed a diverse set of biological processes. Module 1 (n=788) was predominantly enriched for protein kinase activity, histone acetylation, and proteasome-related functions (**S2 Fig & S6 Table**). Module 2 (n=811) was mainly characterized by ribosomal functions and protein translation processes (**S3 Fig. & S7 Table**). Module 4 containing the highest number of genes (n=4781) revealed enrichment in negative regulation of gene expression and nucleic acid metabolism (**S4 Fig & S8 Table**) Modules 5 (n=332) (**S5 Fig & S9 Table**) and Module 8 (n=1147) (**S6 Fig & S10 Table**) displayed similar enrichments related to signal transduction, response to stimulus, and monoatomic ion transport. Module 10 (n=475) exhibited a profile similar to Module 8, featuring enrichments in general transmembrane transport (**S7 Fig & S11 Table**). There were no significantly enriched processes for genes in Module 7 (n=99).

### Identification of core genes that underlie response to ivermectin

Based on the Spearman rank correlation, betweenness and eigenvector centrality emerged as robust indicators for gene significance (**S8 Fig**). Utilizing these metrics, genes within each significant module that ranked in the top 5% were intersected with DEGs, which revealed a total 219 core genes. These core genes were further analyzed to understand their function in response to IVM.

Core genes were divided into nine functional categories (**Table 4 & S12 Table**) using a manual classification delineated in the methods section.

**Table 4:**
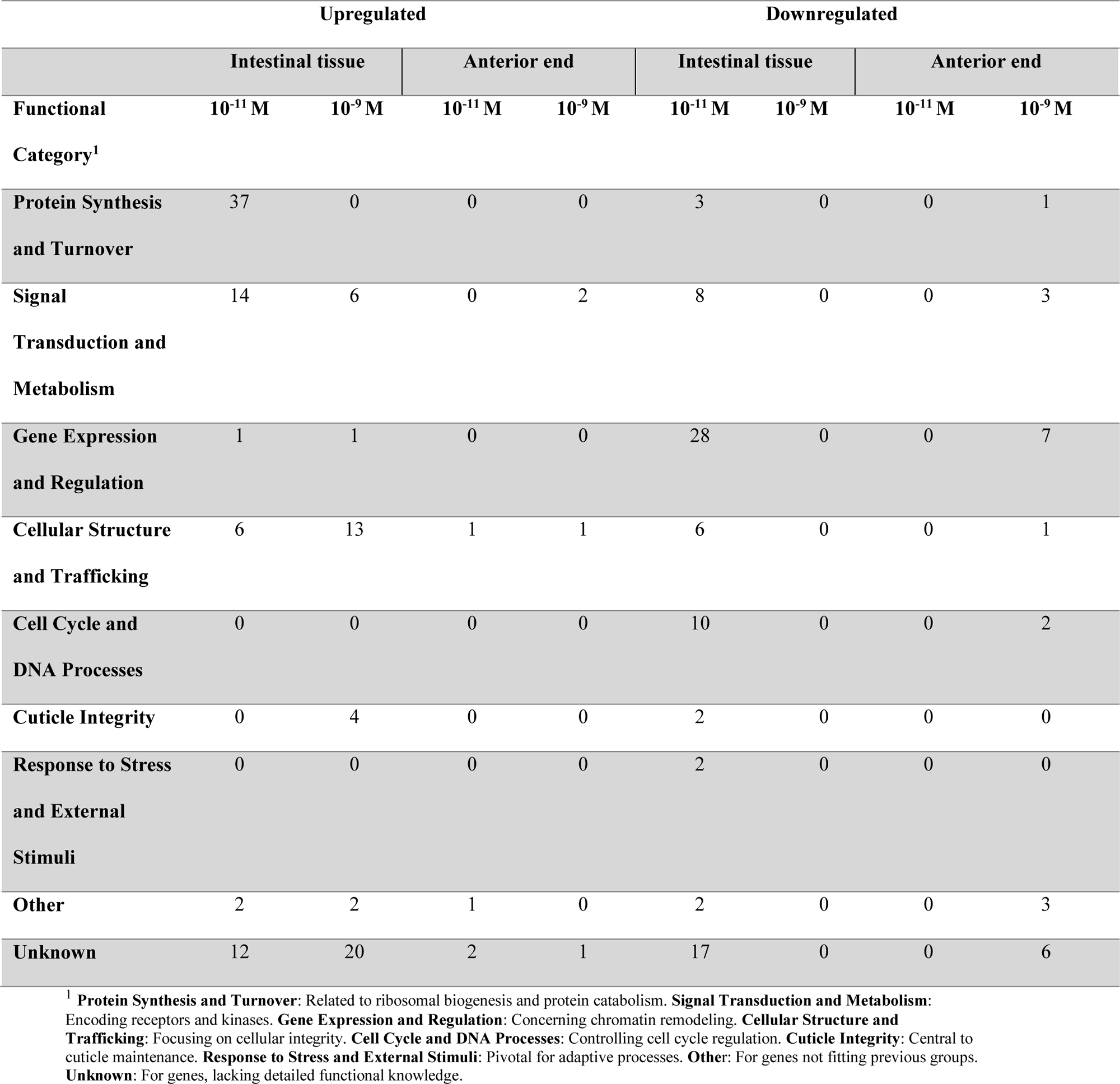
Differential core gene expression in intestinal tissue and anterior end after ivermectin exposure. The table quantifies the number of upregulated and downregulated genes across functional categories in intestinal tissue and anterior end at 10^-11^ M and 10^-9^ M ivermectin concentrations.

### Intestinal tissue core genes

Following exposure to 10^-11^ M IVM, four significant modules (2, 4, 5, and 8) were identified in the intestinal tissue, which resulted in 150 core genes exhibiting several patterns (**Table 5 & S12 Table**). Predominantly, Module 4 exhibited downregulation across multiple functional categories, with a marked emphasis on “Gene Expression and Regulation”, suggesting broad transcriptional arrest. Additionally, a complete downregulation in the “Cell Cycle and DNA Processes” category in the same module indicated an inhibition of cell cycle progression. In contrast, “Signal Transduction and Metabolism” within Module 4 displayed a majority of upregulated core genes, indicating a differential cellular response to IVM. Notably, Module 2 was characterized by exclusive upregulation, dominated by the “Protein Synthesis and Turnover” category. This upregulation was specifically observed in ribosomal genes and select genes involved in protein folding and proteasome activity. Modules 8 and 5 exhibited a more diverse yet restricted expression, with Module 8 showing downregulation across multiple functional categories, and Module 5 containing fewer downregulated genes in “Cell Cycle and DNA Processes” and “Cellular Structure and Trafficking”.

**Table 5:**
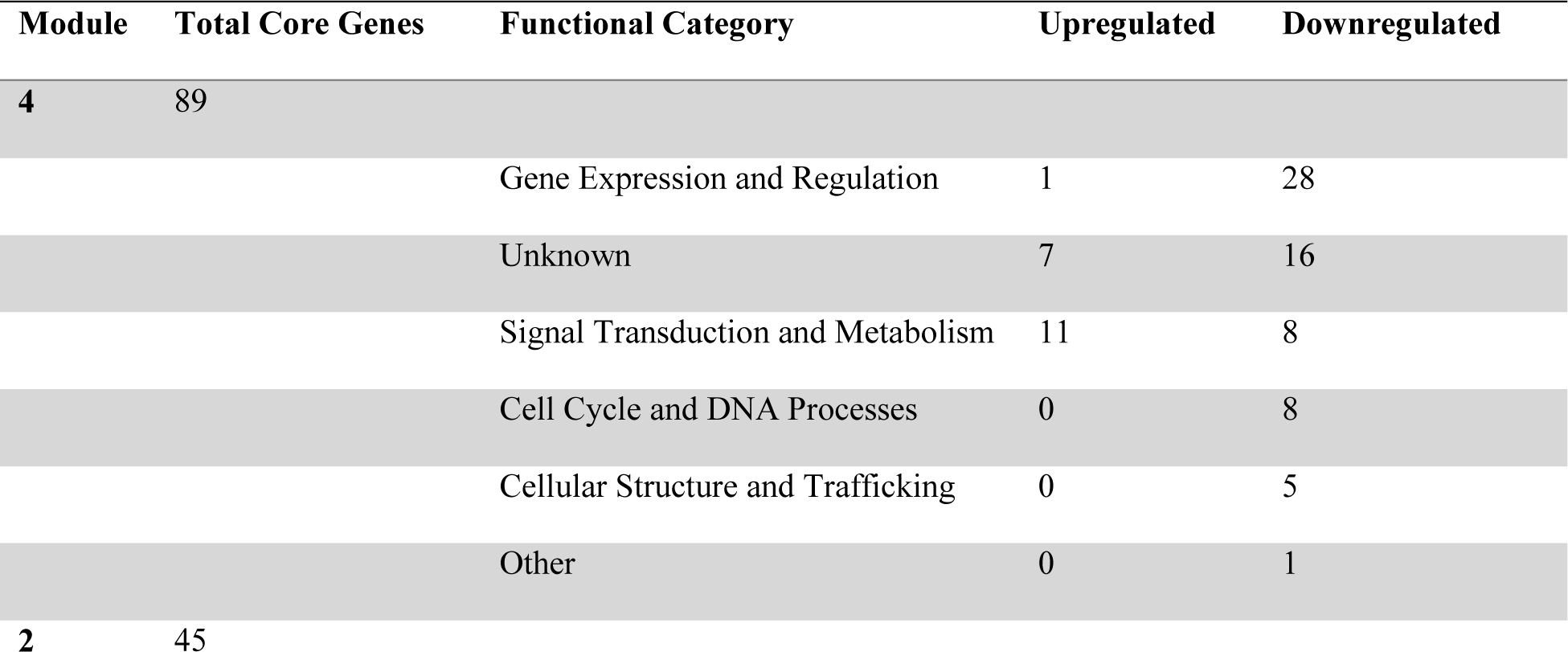
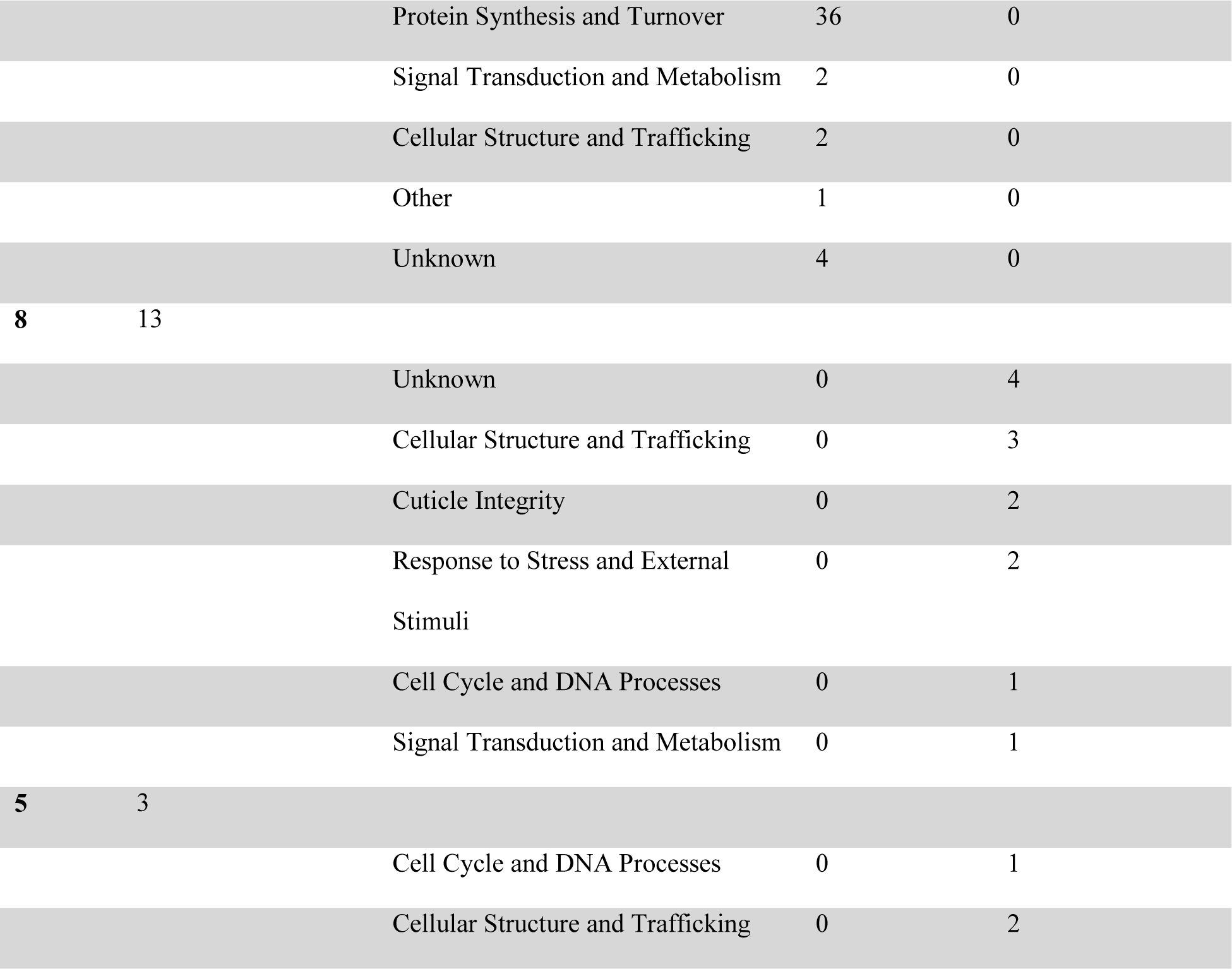
Differential core gene expression in intestinal tissue after exposure to 10^-11^ M ivermectin exposure. The table quantifies the number of upregulated and downregulated genes across functional categories in the intestinal tissue in 10^-11^ M ivermectin concentration.

Three significant modules (1, 5, and 8) were identified after 10^-9^ M IVM exposure (**Table 6 & S12 Table**) consisting of 46 core genes in total. In contrast to findings in 10^-11^ M IVM, an overwhelming 98% (n=45) were upregulated and were integral to various cellular processes. Module 8 mainly displayed upregulated gene expression across various functional categories, most notably in “Cellular Structure and Trafficking”. This suggests a targeted cellular response aimed at maintaining membrane integrity and facilitating intracellular trafficking. Furthermore, the same module showed complete upregulation in both the “Cuticle Integrity” and “Signal Transduction and Metabolism” categories, indicating concerted efforts in cuticle formation and maintenance, as well as the activation of specific intracellular signaling and metabolic pathways. These adaptive responses were recapitulated in Modules 1 and 5.

**Table 6:**
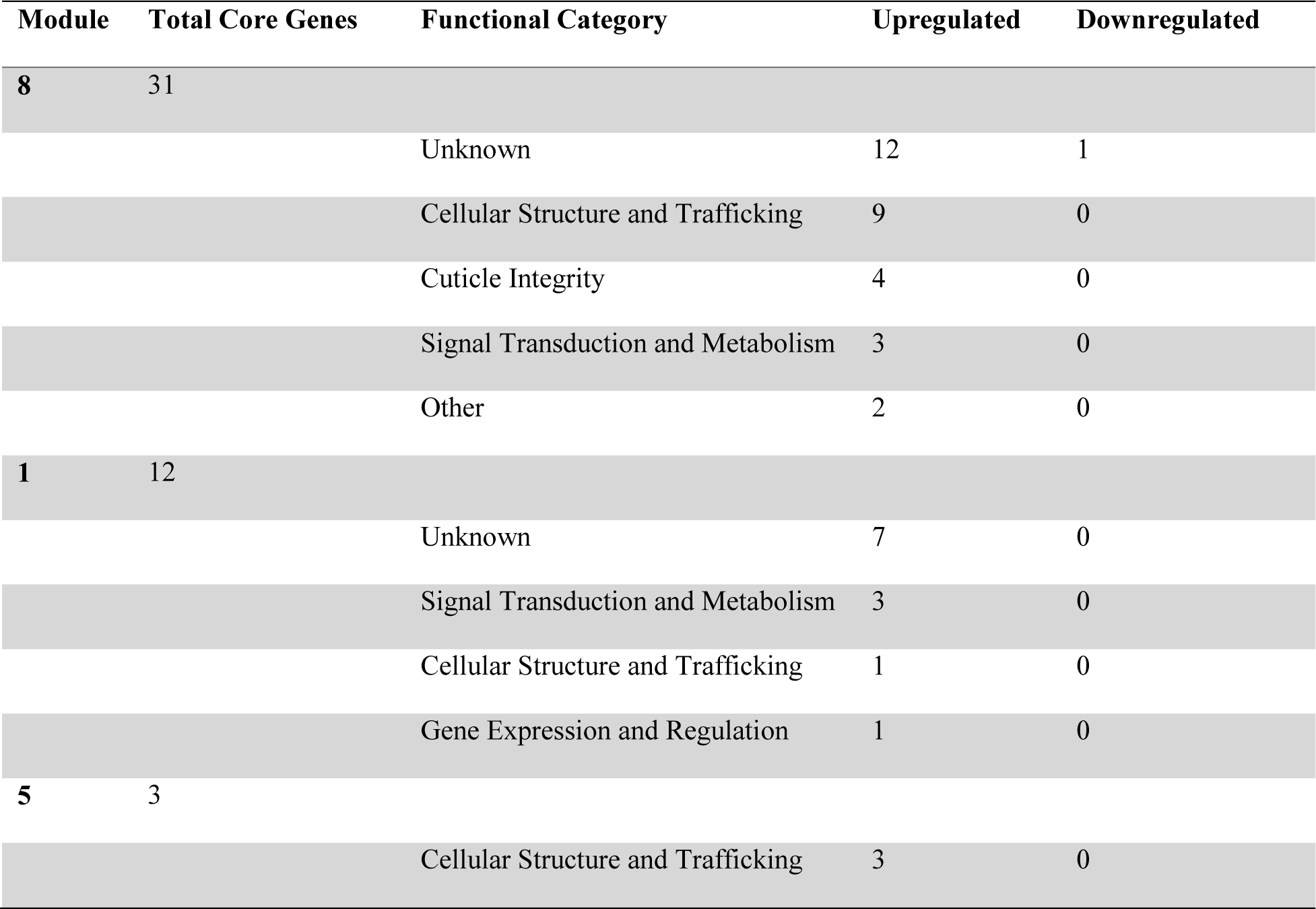
Differential core gene expression in intestinal tissue after exposure to 10^-9^ M ivermectin exposure. The table quantifies the number of upregulated and downregulated genes across functional categories in the intestinal tissue in 10^-9^ M ivermectin concentration.

Overall, the gene expression pattern of core genes was apparent in the intestinal tissue within different IVM concentrations. In 10^-11^ M IVM, most core genes were downregulated, compared to the 10^-9^ M concentration where upregulation was the fundamental response. Interestingly, some core gene exhibited remarkable shifts in expression profiles based on concentration. Specifically, *PgR028_g047 (sorb-1)* transitioned from a 6-fold downregulation at 10^-11^ M to a 5-fold upregulation at 10^-9^ M and belonged to “Cellular Structure and Trafficking” in Module 5. Conversely, *PgB01_g200 (gmap-1)* and *PgR046_g017 (col-37 & col-102)*, which belonged to “Cuticle Integrity” in Module 8, shifted from 15-fold and 37-fold downregulation, respectively, at 10^-11^ M to 4-fold and 3-fold upregulation at 10^-9^ M.

### Anterior end core genes

After exposure to 10^-11^ M IVM, two significant modules (2 and 7) were identified in the anterior end containing four upregulated core genes (**S12 Table**). Module 2 contained one core gene, which belonged to “Cellular Structure and Trafficking” and Module 7 contained three core genes, three categorized to “Unknown”, and one to “Other”.

After exposure to 10^-9^ M IVM, two significant modules, 4 and 10, were identified, comprising 25 core genes (**Table 4 & S12 Table**). Contrasting the findings after exposure to 10^-11^ M IVM, a notable 84% (n=21) of these genes were downregulated, implicating several critical cellular processes in the anterior end. Module 4 was characterized by widespread downregulation, particularly in the “Gene Expression and Regulation” category, suggesting an overarching halt in transcriptional activity. This module also presented a mixed regulation pattern in “Signal Transduction and Metabolism”, potentially indicative of dysregulated intracellular signaling and metabolic pathways. Furthermore, the downregulation observed in the “Cell Cycle and DNA Processes” category postulates disruptions in cell proliferation.

**Table 4:**
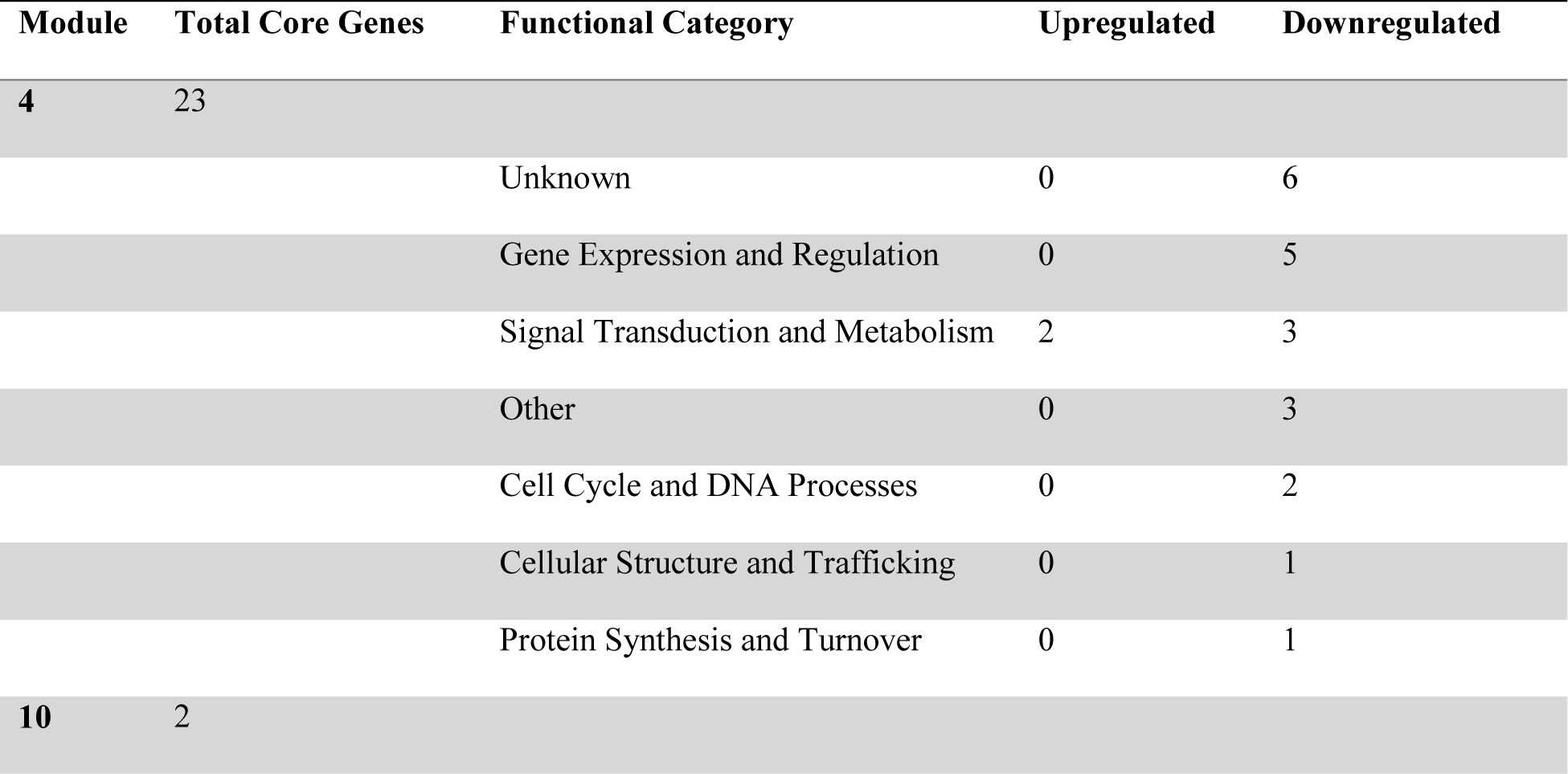
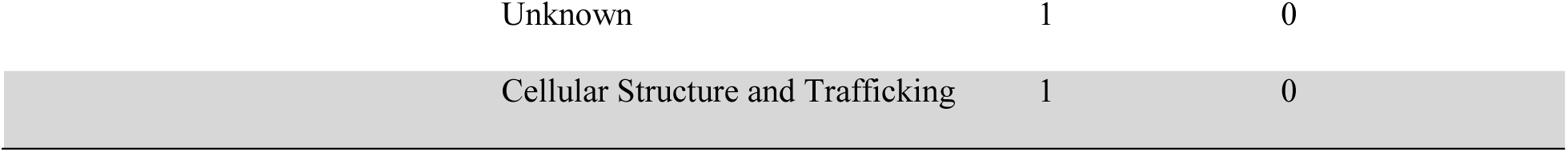
Differential core gene expression in the anterior end after exposure to 10^-9^ M ivermectin exposure. The table quantifies the number of upregulated and downregulated genes across functional categories in the anterior end in 10^-9^ M ivermectin concentration.

| Module | Total Core Genes | Functional Category | Upregulated | Downregulated |
| --- | --- | --- | --- | --- |
| 4 | 23 | Unknown | 0 | 6 |
|  |  | Gene Expression and Regulation | 0 | 5 |
|  |  | Signal Transduction and Metabolism | 2 | 3 |
|  |  | Other | 0 | 3 |
|  |  | Cell Cycle and DNA Processes | 0 | 2 |
|  |  | Cellular Structure and Trafficking | 0 | 1 |
|  |  | Protein Synthesis and Turnover | 0 | 1 |
| 10 | 2 |  |  |  |

|  |  |  |
| --- | --- | --- |
| Unknown | 1 | 0 |
| Cellular Structure and Trafficking | 1 | 0 |

Overall, exposure to IVM induced a substantial difference in the number of core genes between the intestinal tissue and the anterior end. Specifically, the intestinal tissue manifested 190 core genes after IVM exposure, in contrast to the anterior end, which exhibited 7-fold fewer core genes, revealing only 26 genes. Remarkably, only three core genes were shared in between tissue types and were uniformly downregulated. Among these were *PgR223_g002* and *PgR162X_g007* exhibiting an average downregulation of 1.5-fold and 2.6-fold, respectively, and belonged to “Unknown”. Additionally, *PgR050X_g057*, which belonged to “Signal Transduction and Metabolism”, showed an average downregulation of 1.6-fold.

### Functional inference of unknown core genes

We identified 56 core genes of unknown function within six modules (**S13 Table**). To infer their roles, we analyzed the top five upregulated and downregulated core genes based on the ORA of their immediate network neighbors. The upregulated genes, *PgR089_g007, PgR054_g036*, *PgR099_g027*, *PgR169_g001*, and *PgR031_g093*, had neighbors significantly enriched for protein tyrosine phosphatase activity in module 1, indicating potential involvement in signaling pathways. However, *PgR054_g036* from module 8 did not show significant enrichment, suggesting a role in unidentified biological functions. For the downregulated genes, *PgR049_g001* and *PgR024_g021* from module 8 were linked to cell surface processes and defense responses, respectively. Other downregulated genes, *PgR013_g010*, *PgR022X_g052*, and *PgR024_g012*, did not exhibit significant enrichment, hinting at their participation in novel or less-characterized pathways.

### Cross-module connectivity mediated by calcium-signaling gene PgR047_g066

The non-module gene *PgR047_g066 (gegf-1)*, which has a GO term of calcium ion binding, functioned as a pivotal intermodular gene, establishing connections with four distinct gene modules (1, 3, 4 and 9) and interacting with 71 genes (**Fig 4**). To infer the role of *PgR047_g066*, ORA was performed on this gene and its immediate neighbors in the network. This analysis revealed enriched term related to signaling receptor activity (**Fig 4 & S14 Table**), suggesting *PgR047_g066* may have a signaling role. After exposure to 10^-9^ M IVM, this calcium-signaling gene exhibited a notable 12-fold upregulation within the intestinal tissue. Interestingly, while the IVM response was significantly associated with Module 1 and 4, *PgR047_g066* showed the highest number of direct interactions (n=66) within module 9. Of these interacting partners, a mere six genes (*PgB21_g034*, *PgR003_g063*, *PgR010_g006*, *PgR022X_g030*, *PgR032X_g118* and *PgR034_g082*) were upregulated, and two (*PgR003_g108* and *PgR075_g041*) were downregulated.

**Fig 4.**
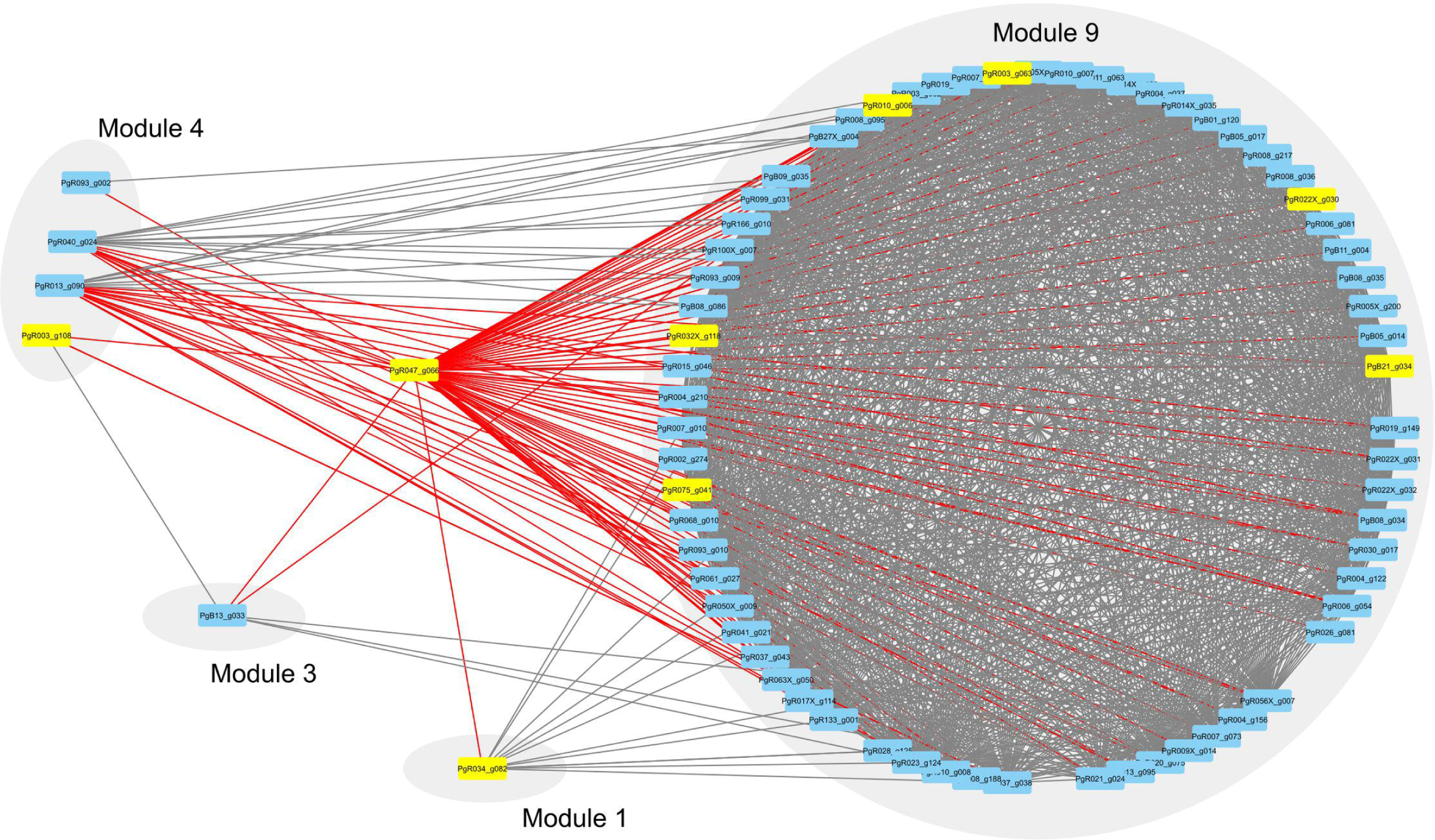
Central role of non-module gene *PgR047_g066* in connecting four gene modules. A gene network showing how *PgR047_g066* serves as a pivotal link among three distinct modules. Rectangles denote individual genes, and edges signify interactions between them. Direct first-degree neighbors to *PgR047_g066* are emphasized with red lines, while differentially expressed genes are colored yellow.

**Fig 5.**
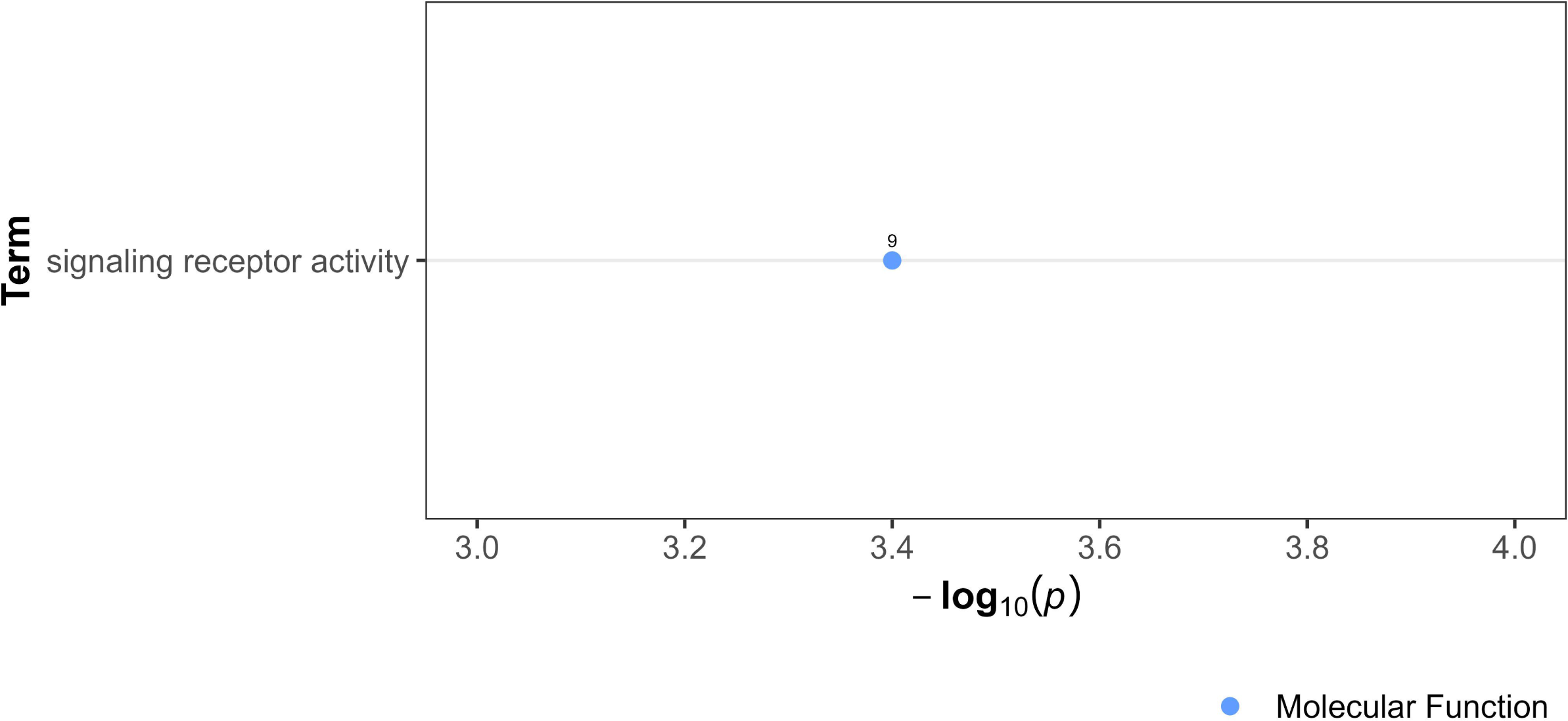
Over-representation analysis showing gene ontology term in *PgR047_g066* and its immediate network neighbors. The data points, color-coded by ontology classification-Biological Process (red), Cellular Component (green), and Molecular Function (blue) are plotted against their respective -log10 transformed p-values, emphasizing the significance of the term. The numerical labels adjacent to each data point indicate the count of differentially expressed genes associated with the respective GO term.

### Interplay between core and candidate genes in response to ivermectin exposure in P. univalens

Prior research on parasitic nematodes and the model organism, *C. elegans*, has proposed a number of candidate genes implicated with ML resistance. These candidate genes were mapped within our gene co-expression network and further analyzed their interaction with the core genes in *P. univalens.* Candidate genes (n= 56) derived from existing literature were selected based on orthologues to *P. univalens* core genes (**S15 Table**). Of these, 43 genes mapped to our network (**S16 Table**), with 21 genes clustering in Module 4. Remarkably, only one candidate gene, *PgR047_g061* (*lgc-37*), encoding a cys-loop GABA ligand-gated chloride channel, was a core gene. Additionally, only eight candidate genes exhibited direct interactions with at least one core gene (**S16 Table**). In Module 4, the candidate genes interacting with core genes were *PgR090_gPgp-10* (*pgp-10*), encoding a P-glycoprotein efflux pump, displayed interactions with two downregulated core genes: *PgR093_g025* (*rrf-1, rrf-2 & ego-1*), an RNA-directed RNA polymerase, and *PgB16_g025* (*C05C8.5*), predicted to possess exonuclease activity. Similarly, candidate gene *PgR025_g110* (*daf-19*), an RFX family transcription factor, interacted with the same two downregulated core genes, suggesting a possible regulatory mechanism. Gene *PgB10_g033* (*mrp-6*), predicted to facilitate ATPase-coupled transmembrane transporter activity, interacted with an upregulated core gene *PgR008_g166* (*mev-1*), which encodes a succinate dehydrogenase cytochrome b560 subunit. The paralogue of *PgB10_g033*, identified as *PgB10_g034* (*mrp-6*), interacted with a downregulated core gene *PgB17_g010* (*F55F10.1*), postulated to play a role in ribosomal large subunit assembly and export. Further, the candidate gene *PgR044_g045* (*dyf-2*), encoding WD-repeat family protein crucial for chemosensation and, interacted with a downregulated core gene of unknown function, *PgR022X_g052*. The candidate gene *PgR053_g053* (*unc-13*), which encodes protein isoforms involved in neurotransmitter release, interacted with another downregulated core gene *PgR028_g087*, predicted to have a function in nucleic acid binding. Lastly, in Module 8, the candidate gene *PgB20_g018* (*eat-2*), encoding a ligand-gated ion channel subunit, interacted with an upregulated core gene *PgR021_g035*, the function of which remains unidentified.

## Discussion

The prevalent resistance to IVM in the equine parasitic nematode, *P. univalens* represents a significant threat to equine health [9, 10] and necessitates an in-depth understanding of its resistance mechanisms. Several previous studies have largely been holistic, examining gene expression across the entire organism [30, 31, 39], which may obscure specific tissue responses critical to resistance in the parasitic worm. Studies that investigate tissue-specific expression often preselect genes, for example intestinal P-glycoproteins [21, 40, 41], potentially missing novel genes of interest. Addressing this, our research conducted an in-depth gene expression analysis in two key anatomical regions: the anterior nerve cell-rich area and the metabolically active intestine. We report a marked contrast in IVM-induced transcriptional responses between these tissues, with notably higher transcriptional activity in the intestine. Furthermore, we utilize a gene network analysis to reduce bias inherent in candidate gene preselection. This method led to the identification of 219 core genes associated with IVM response, offering new insights into potential resistance mechanisms in *P. univalens*.

Upon exposure to 10^-11^ M IVM, no enriched biological processes were seen in the anterior end compared to the intestinal tissue. In the intestinal tissue, transcriptional response was characterized with 58% downregulated genes. Such genes were involved in regulation of cellular processes such transcription and signal transduction, possibly reflecting a resource-conserving stress response. Concurrently, upregulated genes in the intestine after 10^-11^ M IVM suggested enhanced peptide metabolism, ribosomal functions, and energy production, possibly a defense against IVM toxicity. These biological processes in the upregulated genes have also been reported to be enriched in DEGs from other IVM exposed parasitic nematodes, like *H. contortus* larvae [31] and adult *Brugia malayi* [30], implying a shared stress response strategy across species. After exposure to 10^-9^ M IVM, the intestinal tissue exhibited a reversal in gene expression patterns, predominantly characterized by upregulation. This gene expression shift was particularly notable in genes associated with ion transport, cuticle structural integrity, and protein tyrosine phosphatase activity. The nematode cuticle, a critical defensive layer against environmental assaults [42], including anthelmintic agents, adapts by altering its permeability [43]. This adaptation presumably mitigates drug efficacy, with the cuticle also considered to be a principal site for drug uptake [25]. Consequently, the observed upregulation in genes governing cuticle maintenance and structure likely represents a strategic, protective response to IVM exposure. The role of protein tyrosine phosphatases is orchestrating a myriad of cellular processes, such as signal transduction and cell cycle regulation [44]. Therefore, the marked upregulation of genes involved in protein tyrosine phosphatase activity post-IVM exposure highlights a potentially critical adaptive mechanism, possibly aimed at preserving cellular integrity and function. However, the anterior end exhibited a different pattern of increased expression characterized by genes related to proton transport and ATPase activity, may suggest compensatory energy production to maintain neural function under IVM exposure. Overall, this reveals *P. univalens* transcriptional responses to IVM between tissues types, with distinct adaptations in each, suggesting an organism-level strategy for mitigating stress to xenobiotic exposure. This comprehensive difference in response between the anterior end and the intestine highlight the complexity of anthelmintic exposure.

From the broader transcriptional patterns and responses to a more focused analysis, the study identified 219 core genes predominantly expressed in the intestinal tissue, encompassing diverse functional categories. While only four core genes were identified in the anterior end at 10^-11^ M IVM concentration, their biological pattern remained unclear. In contrast, in the intestine at the same IVM concentration, there was a clear pattern of transcriptional arrest and reduced cell proliferation, aligning with the downregulation of core genes associated with these processes. Similarly, in a study of *B. malayi*, gene expression changes related to meiosis were significantly downregulated 24 hours after IVM exposure [30]. This downregulation included genes whose RNAi phenotypes in *C. elegans* lead to developmental arrests [30]. A similar set of core genes such as, *PgB03_g135* (*wapl-1*), *PgR024_g010* (*capg-2*) and *PgR027_g140* (*tln-1*) were seen in the current study, suggesting a potentially conserved response mechanism across different nematode species to IVM exposure. Concurrently, were observed upregulation of core genes involved in protein synthesis and management of misfolded/unfolded proteins, congruent with the ORA results in this study and other studies [30, 31, 45, 46]. Interestingly, enrichment of these protein synthesis processes was also reported in unexposed IVM-susceptible and resistant *H. contortus* [31], suggesting that IVM exposure increases the magnitude of expression due to detoxification. Increasing the IVM concentration to 10^-9^ M induced notable shifts in the expression patterns of core genes in both tissue types. In the intestinal tissue, this elevation resulted in the upregulation of core genes implicated in maintaining cellular integrity, signal transduction, metabolism, and cuticle integrity. Interestingly, the anterior end exhibited similar biological processes as the intestinal tissue at the lower IVM concentration (10^-11^ M), particularly with respect to transcriptional suppression and the inhibition of cell division. We identified core genes using eigenvector centrality and betweenness metrics. Eigenvector centrality assesses a gene’s influence based on its connections, while betweenness considers its role as a critical ‘bridge’ within the network, marking these genes as central to the network’s function. Typically, transcriptomic studies post-drug exposure in nematodes, including ours, reveal differentially expressed genes related to drug biotransformation, such as cytochrome P450s, UDP-glucuronosyltransferases, and efflux P-glycoproteins [22, 32, 33, 39, 41, 47–49]. Interestingly, none of the core genes in our study was associated with drug biotransformation processes. This deviation indicates a possible shift in biological prioritization towards transcription arrest, energy production, and cell cycle inhibition, rather than the conventional focus on xenobiotic detoxification under toxic stress. Furthermore, three core genes in the intestinal tissue exhibited a notable change of expression from downregulation at the lower concentration to upregulation at the higher concentration. This includes the gene *PgR028_g047 (sorb-1)*, which plays a role in muscle organization in *C. elegans* [99] and suggests its significance in modulating muscle physiology under anthelmintic stress, potentially contributing to resilience or resistance. Similarly, cuticular genes such as *PgB01_g200 (gmap-1)* and *PgR046_g017 (col-37 & col-102)* followed this trend. Given that *PgB01_g200 (gmap-1)* encodes a lipid transfer protein, essential for maintaining cuticle permeability [43], emphasizes their potential importance under IVM exposure. Given the close phylogenetic relationship between *P. univalens* and *A. suum* [24], and the proposed predominance of cuticular drug uptake in *A. suum* [25], it is plausible that IVM uptake in *P. univalens* occurs similarly through the cuticle. However, the differential gene expression observed in *P. univalens* raises questions about this prevailing understanding. Notably, the anterior end of *P. univalens*, expected to be rich in GluCls [57, 58] and known for its dominant constitutive expression [50], showed fewer overall DEGs and core genes compared to the intestine. This pattern suggests a more intricate mechanism of IVM absorption, possibly involving both cuticular diffusion and ingestion. Such findings underscore the need to reassess IVM distribution mechanisms in these nematodes to deepen our understanding of anthelmintic resistance dynamics.

The non-module gene *PgR047_g066 (gegf-1)*, with calcium ion binding activity based on GO, emerged as a putative intermodular gene. Its role in signaling was inferred from ORA of its immediate network neighbors, which showed involvement in molecular transducer activity, signaling receptor activity, and transmembrane signaling receptor activity. This finding is particularly noteworthy because calcium signaling plays a crucial role in nematode physiology, especially in muscle function and neurotransmission [51], areas significantly affected by IVM. The 12-fold upregulation of this gene at 10^-9^ M IVM concentration within the intestinal tissue, coupled with its interactions with 71 genes across various modules, points to a potential central role in the nematode’s response to IVM. This makes *PgR047_g066 (gegf-1)* a promising candidate for further research to understand its impact in IVM response.

We compared our core genes with known resistance genes (candidate genes) to understand IVM resistance in *P. univalens*. Of the 45 candidates mapped to the network, only the cys-loop GABA ligand-gated ion channel gene PgR047_g061 (lgc-37) emerged as a core gene and was found to be downregulated in the intestinal tissue. This is in contrast to a previous study that reported upregulation of *lgc-37* in the anterior end post-IVM exposure[22]. The difference in findings can be attributed to our application of median filtering in DESeq2 [52], which is a more conservative approach than the mean filtering used in earlier studies [22, 32]. When mean filtering was applied in our current study (results not shown), we observed a significant upregulation of *lgc-37* in the anterior end at a concentration of 10^-9^ M IVM. The observed expression of *lgc-37* in both tissue types is consistent with the known distribution of cys-loop GABA channels in *C. elegans*, which are predominantly expressed in the nerve-rich anterior end but also expressed in the intestinal tissue [53]. In addition, our findings align with other research emphasizing the importance of ligand-gated ion channel genes, like lgc-54 in *Teladorsagia circumcincta* [54] and *lgc-26* in *C. elegans* [33], in nematode IVM response and resistance.

In summary, this study provides novel insights into the tissue-specific and dose-dependent responses of *P. univalens* to IVM. Through an unbiased gene network approach, the study identified 219 core genes, unraveling the complex interplay of gene interactions and processes governing drug response and possibly resistance. Furthermore, by conducting ORA on the immediate neighbors of the unknown core genes, we have made strides in inferring their possible functions. The unique responses illuminated by the core genes offer a deeper understanding of the parasite’s key mechanisms for coping with different IVM concentrations. This level of insight, unattainable through ORA alone, showcases the utility of the gene network approach in dissecting complex biological responses. Prospective research should functionally interrogate the role of these core genes in IVM response and resistance. Furthermore, augmenting this approach with comprehensive proteomic and metabolomic profiles across larger sample sizes of IVM-susceptible and -resistant worms could significantly advance the understanding of the drug’s response and resistance mechanisms.

## Materials and methods

### Data source and ivermectin treatment across two experiments

In this study, we used a comprehensive array of gene co-expression network inference algorithms, facilitated by the Seidr toolkit [37], to delineate the core genes associated with IVM response in *P. univalens*. Acknowledging that increased sample sizes strengthen both the statistical robustness and the generalizability of network inferences [38, 55], we aggregated RNA-sequencing data derived from the anterior end and intestine across 28 individual *P. univalens* adult worms. These worms were collected from two independent experiments conducted in 2017 [22] and 2019 [34]. Briefly, alive adult *P. univalens* worms were harvested from anthelmintic-naïve Icelandic foals at Selfoss abattoir and *in vitro* exposed to 10^-11^ M (n=6), 10^-9^ M (n=12) and control, without the addition of IVM (n=10) for 24 h. After incubation, the worms were dissected and the anterior end and the intestinal tissue preserved in RNAlater.

### RNA sequencing

Worms from both experiments underwent a standardized RNA extraction and sequencing workflow. Anatomical segments of the worms, specifically the anterior end and the intestine, were processed individually. Approximately one microgram of RNA from each tissue sample was utilized to prepare libraries using the TruSeq Stranded mRNA Library Preparation Kit with polyA selection (Illumina Inc., San Diego, USA). The libraries were paired-end sequenced in two separate batches, A (9 worms) [22] and B (19 worms). Sequencing batch A underwent sequencing for 200 cycles on an S1 flow cell, and B for 300 cycles on an S4 flow cell. All sequencing procedures were conducted on the Illumina NovaSeq 6000 platform by the National Genomics Infrastructure at Science for Life Laboratory, Sweden.

### Read processing, mapping and quantification

In the downstream analysis, we integrated sequencing data derived from both experiments and sequencing batches. Any variations due to these contrasting sources of data was statistically controlled for during model building. For simplicity, the anterior part of the worm is referred to as “anterior end”, while the intestine is called “intestinal tissue”. These terms are collectively referred to as “tissue types” throughout the manuscript.

Ribosomal RNA reads were evaluated and filtered using SortMeRNA (v4.3.6) and the smr_v4.3_fast_db rRNA database [56]. To perform quality trimming and adapter removal, Trimmomatic (v0.39) [57] was used with the following customized parameters: a sliding window of length four, a minimum quality threshold of 20, and a minimum length requirement of 36. FastQC (v0.11.9) [58] was then applied to assess the quality of the obtained reads.

Salmon (v1.9.0) [59] was employed for expression quantification against the *P. univalens* transcriptome, utilizing the full genome (PRJNA386823, release WBPS17 (WS282)) as the decoy sequence, which is accessible at WormBase Parasite [60]. The RNA sequencing raw files can be found at the ENA database via at https://www.ebi.ac.uk/ena/browser/view/ (accession numbers: PRJEB37010 and PRJEB70467 for sequencing batch A and B, respectively). Moreover, the analysis pipeline code and necessary files are available at https://github.com/ruqse/Parascaris-IVM-GeneNetwork.

All RNA-based analyses were conducted using R [61] version 4.2.2. The Salmon output was converted into a matrix with the tximport R package (v1.22.0) [62], incorporating annotations related to the PRJNA386823 WBPS17 release available at WormBase Parasite [60] for subsequent transcript/gene-level analysis.

### Differential gene expression

Differential gene expression analysis was performed on raw count data using the DESeq function in the DESeq2 R package [52], with the following model design:

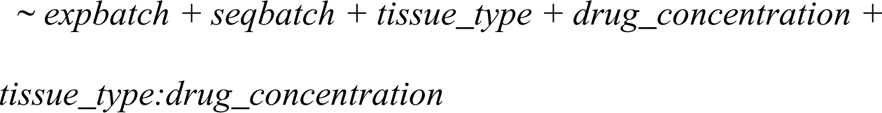

This model accounted for experiment batch, sequencing batch, tissue type, IVM concentration, and the interaction between tissue type and concentration. The introduction of batch variables into the model was a strategic step aimed at mitigating any potential biases arising from batch effects. Differentially expressed genes (DEGs) were identified based on an adjusted *p*-value < 0.05 and a Log2foldchange (Log2FC) ≥ 0.5 or ≤ -0.5 (equivalent to a 1.4-fold change), adhering to the guidelines proposed by Schurch et al. [63]. A filter was implemented to further refine the analysis, considering the median expression value of each gene across all samples.

### Over-representation analysis

Over-representation analysis (ORA) was conducted on DEGs using a background of 12032 genes with non-zero read counts across all samples. The analysis utilized the Gene Ontology (GO) database (Release 2023-07-27) and was implemented via the *gost* function from the gprofiler2 R package (v0.2.2) [64], setting *P. univalens* (ASM225920v1 (2020-05-WormBase)) organism of interest. The highlight parameter was set to TRUE to identify “driver” gene sets in GO results, streamlining data interpretation by clustering similar terms and selecting the most informative. While all significant terms were tabulated, for easier visualization, only “driver” GO terms were plotted.

### Gene network construction, inference and visualization

For this analysis, we only considered genes that exhibited non-zero total read counts. To identify highly correlated genes that potentially share similar biological functions, pathways, or are co-regulated, we performed gene co-expression network analysis using the Seidr tool kit [37] on count data. Ten different gene network inference methods, namely ARACNE, CLR, GENIE3, Ensemble method (llr), NARROMI, Partial correlation, Pearson, PLSNET, Spearman, and Tomsimilarity, were to the processing of building meta networks utilizing the Seidr toolkit. The individual networks obtained from each inference method were aggregated into a consensus network using the “Seidr aggregate” subprogram employing the inverse rank product (IRP) algorithm [65]. However, consensus networks often exhibit dense and spurious connectivity, which can introduce noise and complexity. To address this, backboning was employed to enhance the interpretability of the consensus network by reducing noise and revealing the underlying structure and important gene relationships. Backboning was performed using a range of thresholds: 2.33, 2.05, 1.88, 1.75, 1.64, 1.55, 1.48, 1.41, 1.34, and 1.28. By applying these thresholds, the backboning process effectively reduced noise and retained the most significant connections in the consensus network, allowing for a clearer interpretation of the gene relationships. A gold standard network was built to facilitate selection of the most optimum backboned consensus network. To build the standard network, *C. elegans* genes that have orthologues in *P. univalens* and their respective KEGG pathway annotations taken from the AnnotationDbi R package [66] were used. The “Seidr roc” subprogram was used to calculate true positive rate and false positive rate and derive the corresponding area under the curve (AUC), as a measure of accuracy. The backboned consensus network with the highest AUC, 0.68, which corresponded to a threshold of 1.34, was selected for downstream analyses.

A community detection tool, Leiden [67], as implemented in the igraph R package [68] was applied to detect co-expression modules and respective sub-networks within the consensus network. Our application of this tool adhered to the default parameters, with the exception of the resolution and the number of algorithm iterations, which were specifically set at 1 and 1000, respectively.

We employed a hypergeometric test to identify modules significantly linked to differential expression after IVM exposure. This test assessed the overlap between genes in each module and the differentially expressed genes. Modules with a *p*-value less than 0.05 were chosen for further analyses. Gene set enrichment analysis was then conducted on these pertinent modules, shedding light on the inherent biological processes delineated by the module detection.

To identify the most pivotal genes in each significantly associated module, we employed various centrality metrics including Eigenvector, Betweenness, Closeness, and Pagerank. To mitigate redundancies, a correlation heatmap facilitated the selection of the least correlated metric(s), which was then utilized to rank genes. Genes that overlapped with DEGs and ranked in the top 5% (95th percentile) for each module were classified as core genes, warranting further analysis. To streamline the analysis, core genes were systematically categorized in groups using a multifaceted approach. Criteria incorporated GO annotations, established biological functions, outcomes from BLAST searches, and data pertinent to their *C. elegans* orthologues. The latter encompassed GO classifications, phenotypes discerned via RNA interference (RNAi), allele-specific phenotypes, and succinct descriptions sourced from WormBase. For core genes with unidentified functions, inferences were drawn based on the functions or biological processes associated with their immediate network neighbors.

To refine the identification of key genes, we employed two distinct approaches. First, we identified DEGs that did not belong to any specific module but served as connectors between multiple modules. Second, we selected candidate genes known to be associated with ML resistance, mapped them onto the network, and assessed whether their immediate neighbors were classified as core genes.

## Supporting information

**S1 Table. Differentially expressed genes in the anterior end and intestinal tissue of *Parascaris univalens* exposed to 10^-11^ and 10^-9^ M ivermectin.**

**S2A Table. Over-representation analysis of downregulated genes in the intestinal tissue of *Parascaris univalens* exposed to 10^-11^ M ivermectin.**

**S2B Table. Over-representation analysis of upregulated genes in the intestinal tissue of *Parascaris univalens* exposed to 10^-11^ M ivermectin.**

**S3 Table. Over-representation analysis of upregulated genes in the intestinal tissue of *Parascaris univalens* exposed to 10^-9^ M ivermectin.**

**S4 Table. Over-representation analysis of upregulated genes in the anterior end of *Parascaris univalens* exposed to 10^-9^ M ivermectin.**

**S5 Table. Overlap statistics showing seven gene modules significantly associated with ivermectin response.**

**S6 Table. Over-representation analysis of Module 1 genes.**

**S7 Table. Over-representation analysis of Module 2 genes. S8 Table. Over-representation analysis of Module 4 genes.**

**S9 Table. Over-representation analysis of Module 5 genes. S10 Table. Over-representation analysis of Module 8 genes.**

**S11 Table. Over-representation analysis of Module 10 genes.**

**S12 Table. Differentially expressed core genes across seven significant modules in the anterior end and intestinal tissue of *Parascaris univalens* exposed to 10^-11^ and 10^-9^ M ivermectin.**

**S13 Table. Differentially expressed core genes of unknown function in the anterior end and intestinal tissue of *Parascaris univalens* exposed to 10^-11^ and 10^-9^ M ivermectin.**

**S14 Table. Over-representation analysis of non-module *PgR047_g066* and its immediate neighbors.**

**S15 Table. Literature-derived candidates genes implicated in resistance to Macrocylic lactone drug class.**

**S16 Table. Literature-derived candidates genes implicated in resistance to Macrocylic lactone drug class that mapped to the consensus network.**

**S1 Fig. Overlap statistics showing seven gene modules significantly associated with ivermectin response.** The bar charts represent the percentage overlap between differentially expressed genes (DEGs) and gene modules, illustrated for both 10^-9^ M and 10^-11^ M concentrations. Blue bars indicate the percentage of DEGs within each module, while red bars show the percentage of each module’s genes that are differentially expressed. Statistical significance is marked by asterisks above the bars (\*\*\**p*<0.001, \*\**p*<0.01).

**S2 Fig. Over-representation analysis showing gene ontology terms in of Module 1 genes.** The data points, color-coded by ontology classification-Biological Process (red), Cellular Component (green), and Molecular Function (blue) are plotted against their respective -log10 transformed p-values, emphasizing the significance of each term. The numerical labels adjacent to each data point indicate the count of genes.

**S3 Fig. Over-representation analysis showing gene ontology terms in of Module 2 genes.** The data points, color-coded by ontology classification-Biological Process (red), Cellular Component (green), and Molecular Function (blue) are plotted against their respective -log10 transformed p-values, emphasizing the significance of each term. The numerical labels adjacent to each data point indicate the count of genes.

**S4 Fig. Over-representation analysis showing gene ontology terms in of Module 4 genes.** The data points, color-coded by ontology classification-Biological Process (red), Cellular Component (green), and Molecular Function (blue) are plotted against their respective -log10 transformed p-values, emphasizing the significance of each term. The numerical labels adjacent to each data point indicate the count of genes.

**S4 Fig. Over-representation analysis showing gene ontology terms in of Module 4 genes.** The data points, color-coded by ontology classification-Biological Process (red), Cellular Component (green), and Molecular Function (blue) are plotted against their respective -log10 transformed p-values, emphasizing the significance of each term. The numerical labels adjacent to each data point indicate the count of genes.

**S5 Fig. Over-representation analysis showing gene ontology terms in of Module 5 genes.** The data points, color-coded by ontology classification-Biological Process (red), Cellular Component (green), and Molecular Function (blue) are plotted against their respective -log10 transformed p-values, emphasizing the significance of each term. The numerical labels adjacent to each data point indicate the count of genes.

**S6 Fig. Over-representation analysis showing gene ontology terms in of Module 8 genes.** The data points, color-coded by ontology classification-Biological Process (red), Cellular Component (green), and Molecular Function (blue) are plotted against their respective -log10 transformed p-values, emphasizing the significance of each term. The numerical labels adjacent to each data point indicate the count of genes.

**S7 Fig. Over-representation analysis showing gene ontology terms in of Module 10 genes.** The data points, color-coded by ontology classification-Biological Process (red), Cellular Component (green), and Molecular Function (blue) are plotted against their respective -log10 transformed p-values, emphasizing the significance of each term. The numerical labels adjacent to each data point indicate the count of genes.

**S8 Fig. Heatmap and hierarchical clustering of network centrality metrics.** The color-coded matrix displays the Spearman correlation coefficients between different centrality measures, including Eigenvalue, Betweenness, Closeness, Degree, and PageRank, calculated for nodes within the consensus network. Values close to 1 indicate a high positive correlation, illustrated by a gradient from yellow to green, whereas values close to 0 imply no correlation, depicted in purple. The dendrogram reflects the hierarchical clustering based on the correlation values, grouping similar centrality measures.

## Acknowledgments

We thank the National Genomics Infrastructure in Stockholm and Uppsala, funded by the Science for Life Laboratory, the Knut and Alice Wallenberg Foundation, and the Swedish Research Council, for their sequencing services. Our appreciation also extends to the National Academic Infrastructure for Supercomputing in Sweden (NAISS) at UPPMAX, supported by the Swedish Research Council (grant no. 2022-06725), for providing computational resources under projects NAISS 2023/22-119 and NAISS 2023/23-212.

