## Supplementary figures and images for "Gene co-expression network analysis reveal core responsive genes in *Parascaris univalens* tissues following ivermectin exposure"

### S1 Fig

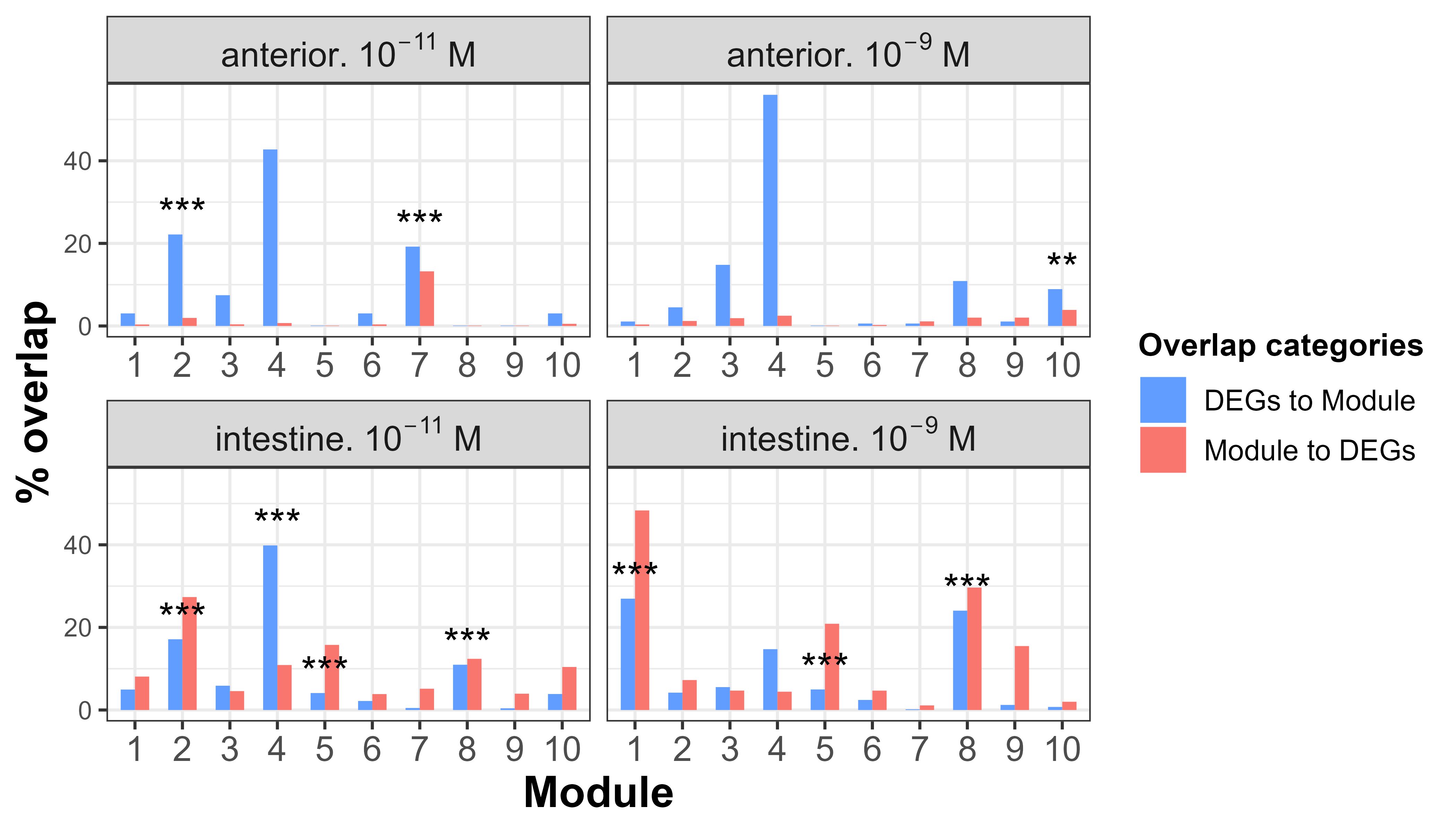

### S2 Fig

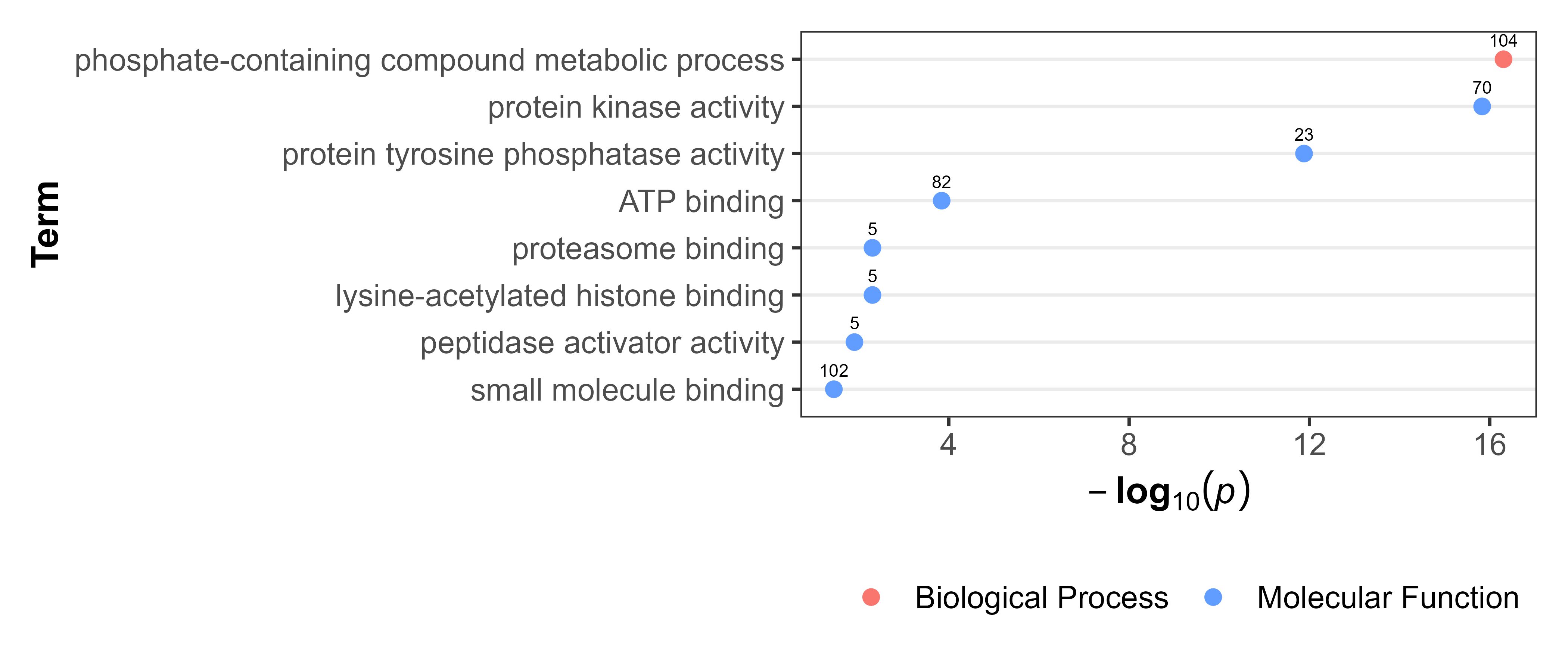

### S3 Fig

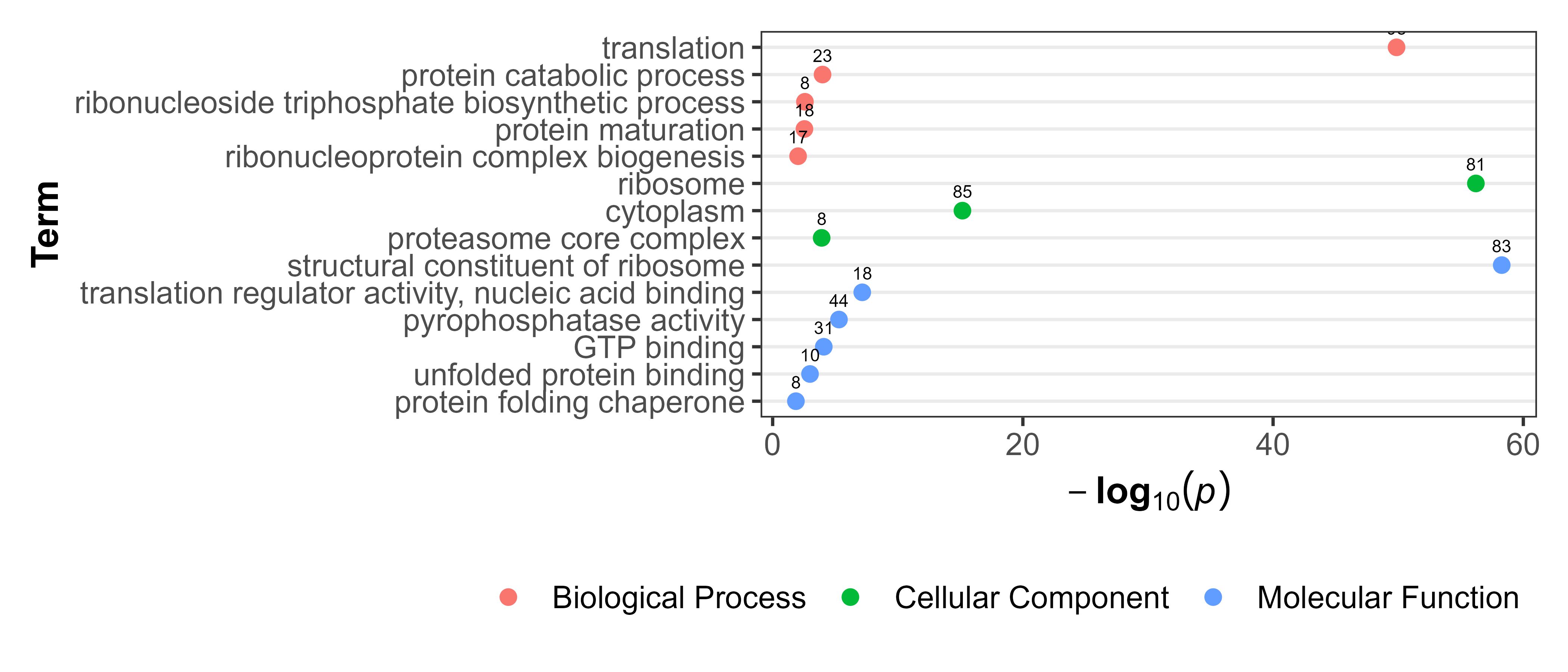

### S4 Fig

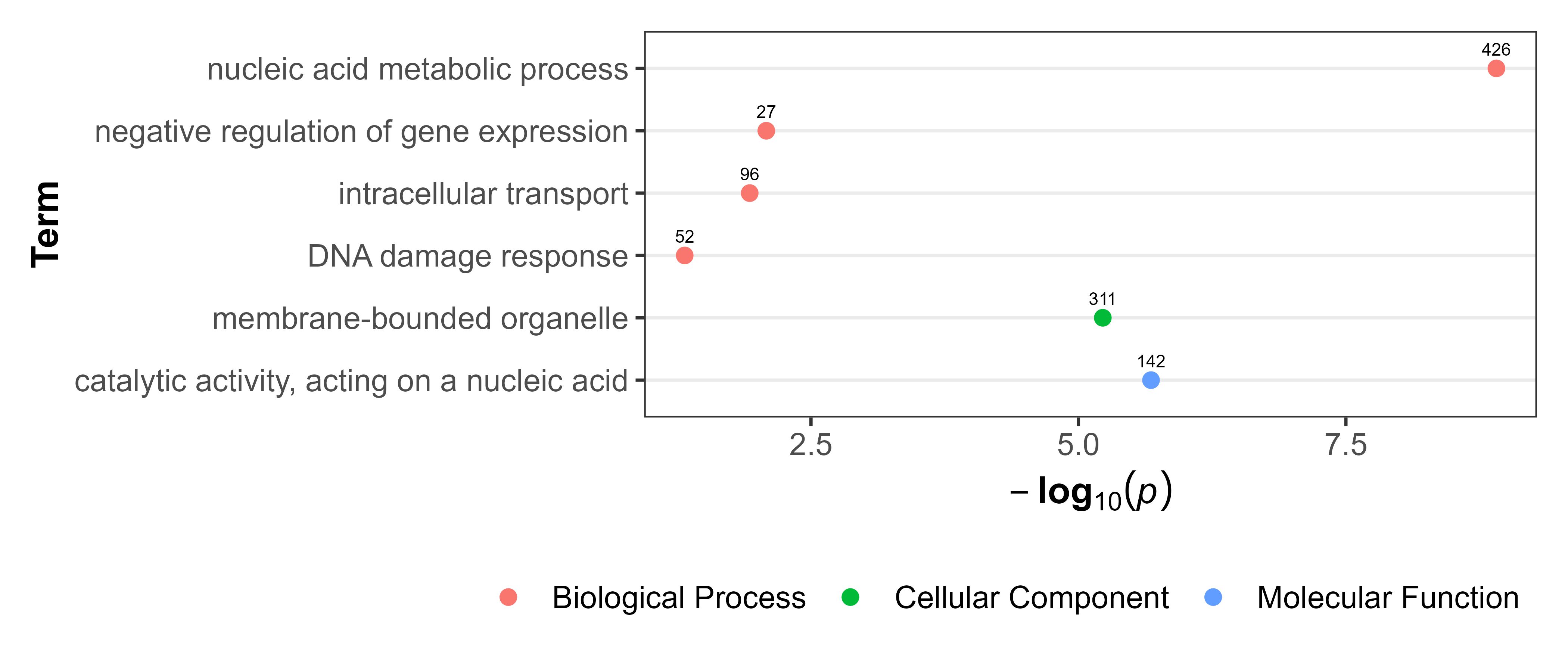

### S5 Fig

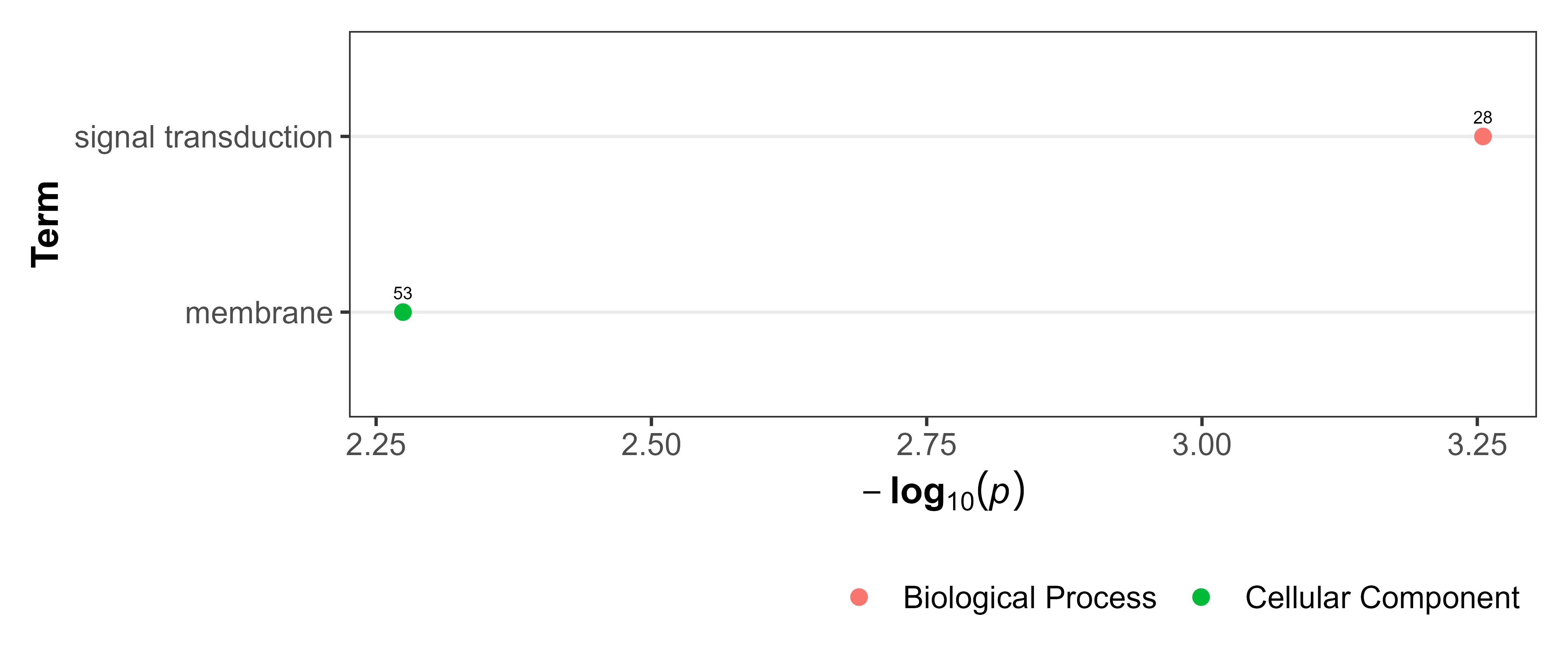

### S6 Fig

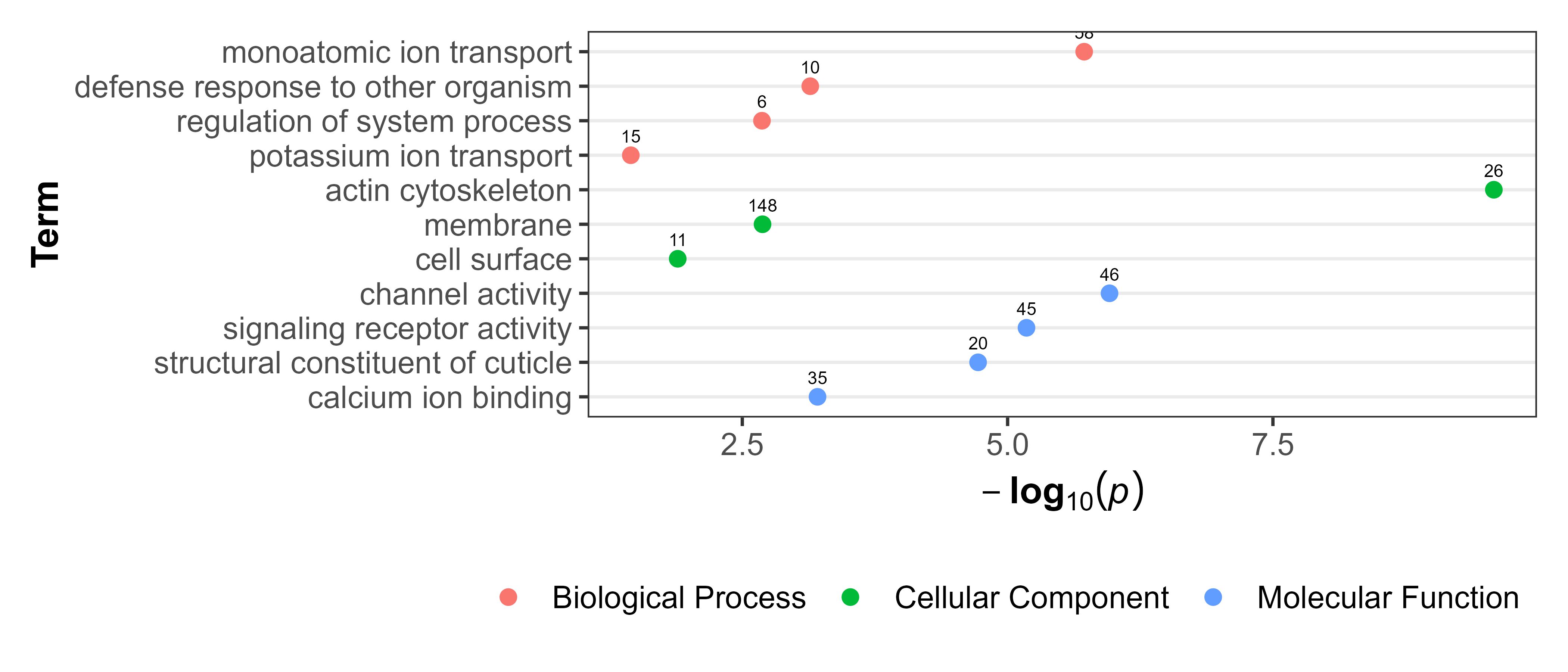

### S7 Fig

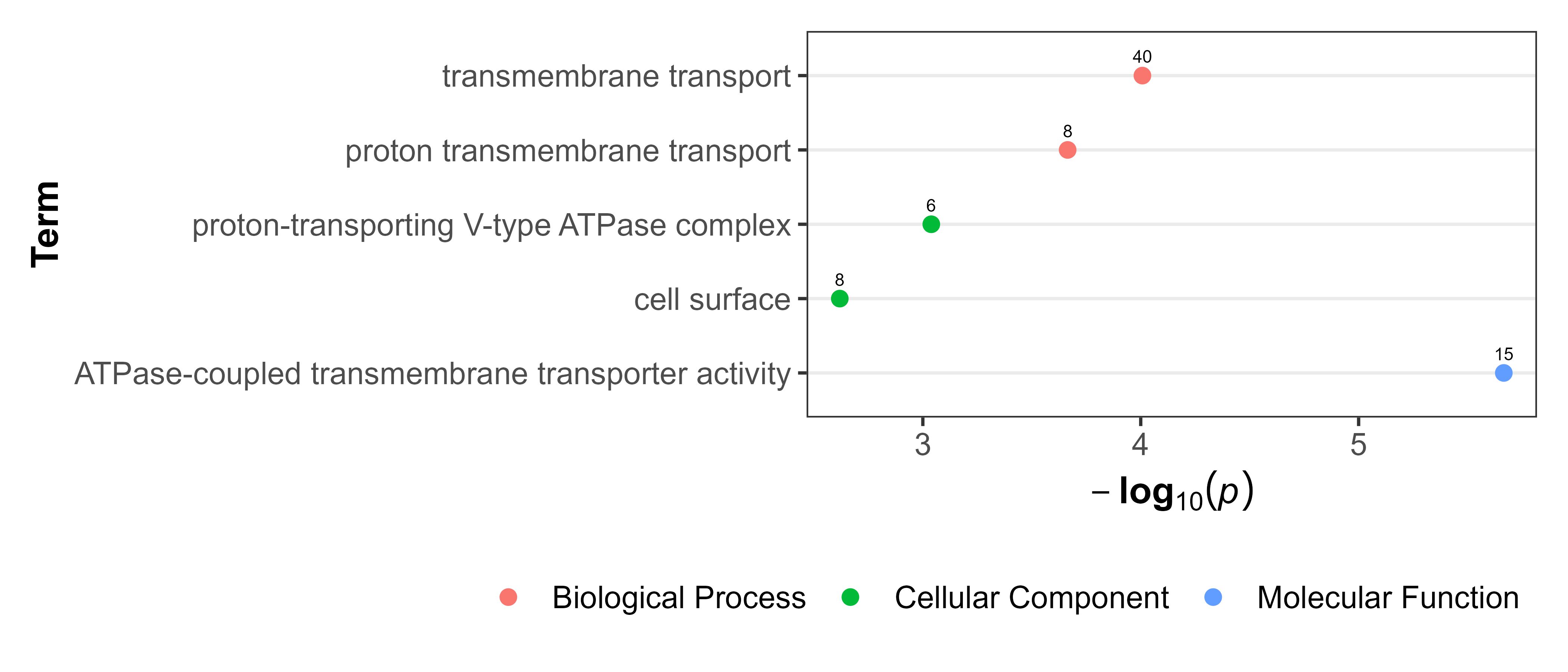

### S8 Fig

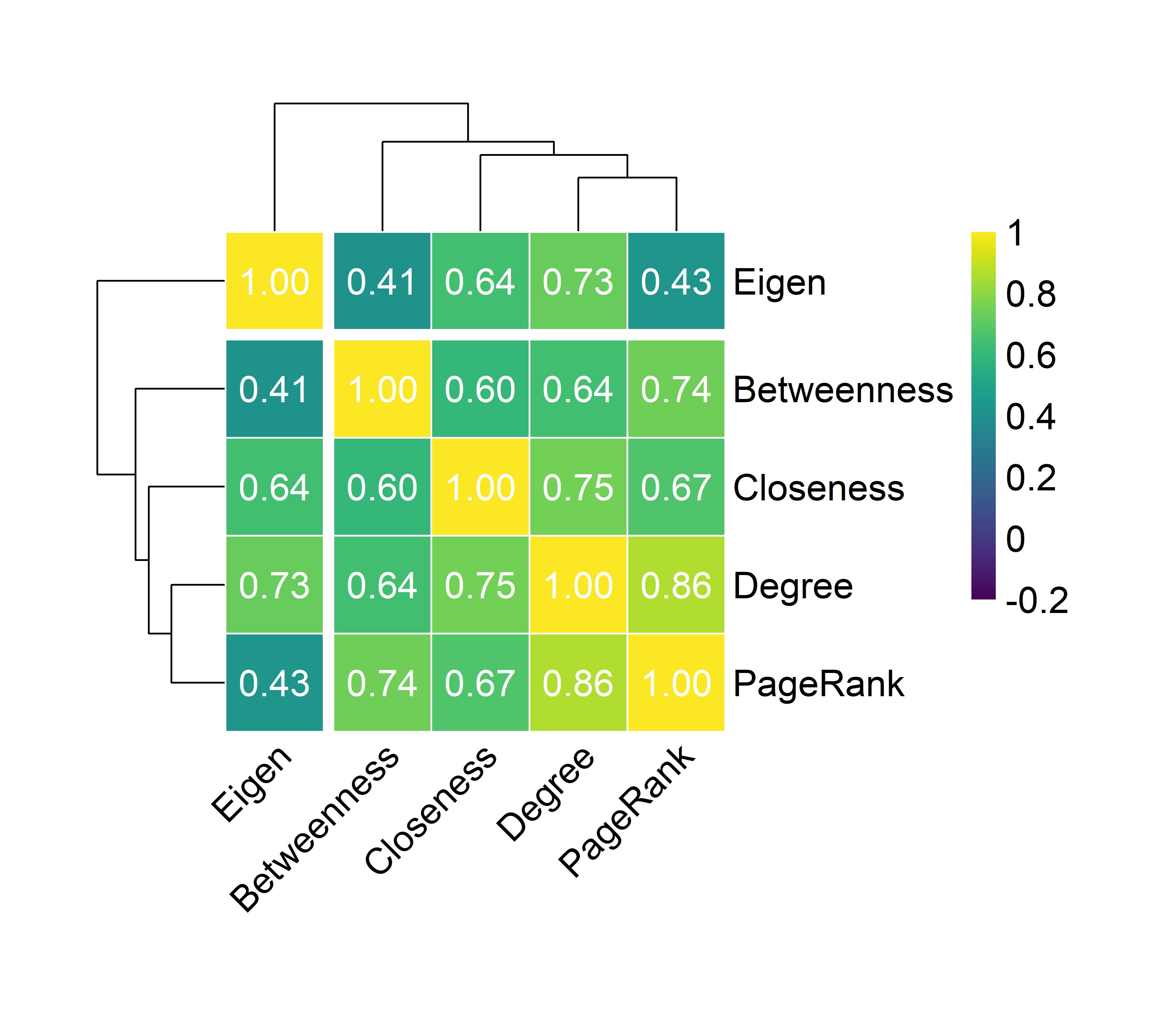
